# Fast and highly efficient affinity enrichment of Azide-A-DSBSO cross-linked peptides

**DOI:** 10.1101/2019.12.17.879601

**Authors:** Manuel Matzinger, Wolfgang Kandioller, Philipp Doppler, Elke H. Heiss, Karl Mechtler

## Abstract

Cross-linking mass spectrometry is an increasingly used, powerful technique to study protein-protein interactions or to provide structural information. Due to sub-stochiometric reaction efficiencies, cross-linked peptides are usually low abundant. This results in challenging data evaluation and the need for an effective enrichment.

Here we describe an improved, easy to implement, one-step method to enrich azide-tagged, acid-cleavable disuccinimidyl bis-sulfoxide (DSBSO) cross-linked peptides using dibenzocyclooctyne (DBCO) coupled Sepharose® beads. We probed this method using recombinant Cas9 and *E. coli* ribosome. For Cas9, the number of detectable cross-links was increased from ~100 before enrichment to 580 cross-links after enrichment. To mimic a cellular lysate, *E. coli* ribosome was spiked into a tryptic HEK background at a ratio of 1:2 – 1:100. The number of detectable unique cross-links maintained high at ~100. The estimated enrichment efficiency was improved by factor 4 −5 (based on XL numbers) compared to enrichment via biotin and streptavidin. We were still able to detect cross-links from 0.25 μg cross-linked *E. coli* ribosome in a background of 100 μg tryptic HEK peptides, indicating a high enrichment sensitivity. In contrast to conventional enrichment techniques, like SEC, the time needed for preparation and MS measurement is significantly reduced.

This robust, fast and selective enrichment method for azide-tagged linkers will contribute to map protein-protein interactions, investigate protein architectures in more depth and help to understand complex biological processes.

## Introduction

Cross-linking mass spectrometry (XL-MS) has emerged as a widely used tool for studying protein-protein interactions and to obtain structural information on protein complexes. It gains increasing importance by providing complementary information to methods such as cryo-electron microscopy, X-ray crystallography analysis or NMR spectroscopy^1–4^. Analysis of XL-MS data, however, remains challenging mainly due to the low abundance of cross-linked peptides. Especially in the field of *in vivo* cross-linking the linker molecule has to permeate the cell membrane and until it has reached reactive amino acid residues in a close enough proximity it is already partly hydrolyzed. This leads to low sub-stochiometric reaction efficiencies.^4–6^ In conclusion, enrichment of cross-linked peptides is crucial. Since cross-linked peptides are on average larger and higher charged, compared to linear peptides, enrichment is often done via size exclusion (SEC)^7–10^ or strong cation exchange chromatography (SCX)^11–13^, respectively. A bottleneck in cross-linking studies regarding complex systems remains, in that coverage is almost exclusively restricted to the most abundant proteins (e.g.^11,14^). To alleviate this issue, cross-linkers with an affinity tag are used, aiming to get a deeper proteome coverage.^6,15–17^ As such e.g. biotin is widely used as affinity tag, due to an effectively working enrichment via streptavidin and the commercial availability of the respective tools.^16–19^

In this study we used azide-tagged, acid-cleavable disuccinimidyl bis-sulfoxide (Azide-A-DSBSO, here termed DSBSO) as published by Kaake et. al.^15^ It is a symmetric, MS cleavable, membrane permeable, homo-bi-functional, N-hydroxysuccinimidyl (NHS) ester-based linker, predominantly reactive with lysine residues. During MS/MS DSBSO generates characteristic doublet ions, thereby circumventing the “n² problem”^20,21^ (the search space increases by n² to the database size). Additionally, DSBSO was shown to be membrane permeable, enabling *in vivo* application.^15^ By that, DSBSO has a very similar chemistry as the previously developed DSSO linker^22^. It additionally contains an azide tag for a selective and bio-orthogonal enrichment using a copper free click reaction^23^ to an alkyne. In the originally published enrichment strategy^15,24^, the cross-linked peptides are first clicked to biarylazacyclooctynone (BARAC) conjugated to biotin, followed by affinity enrichment with streptavidin beads.

Although the Kaake *et al.* have already shown impressive results using DSBSO^15^, we aimed at streamlining the enrichment process, by reducing the number of filtering/working steps to minimize potential sample losses and processing time. Tan *et al.*^17^ have previously reported one step enrichment strategies (based on biotin-avidin affinity) using their one-piece Leiker linker which can be eluted from the enrichment beads by reductive cleavage. To capitalize the advantages of a one step method for DSBSO (or in theory any other azide tagged molecule) we directly enriched cross-linked peptides on alkyne functionalized beads in conjunction with a similar copper-free click reaction. By omitting the use of biotin, we additionally circumvent a potential co-enrichment of endogenously biotinylated proteins. The recovery of the presented method is higher, and the protocol leads to significantly increased final cross-link numbers, when compared to the previous method. In conclusion we show that it is a very valuable tool for future cross-linking studies on complex biological samples, such as tissues or cellular material.

## Materials and Methods

### Materials

Purified *E. coli* ribosome (*E. coli* B strain) was purchased from New England Biolabs (MA, USA) and diluted with dilution buffer (50 mM HEPES, 50 mM KCl, pH 7.5, 10 mM MgAc_2_) to a concentration of 1 mg/mL. Purified recombinant Cas9 from S. Pyogenes fused with a Halo-tag was generated in house, as described by Deng *et al.*^25^ DSBSO was synthesized similar as described by Burke et al^24^. For enrichment similar as described by Kaake *et al.*^15^ (BARAC Method), Dibenzocyclooctyne-PEG4-biotin conjugate (#760749-5MG, Sigma-Aldrich) was used without further purification. Biotin was pulled using Pierce™ High Capacity Streptavidin Resin (# 20359, Thermo). DBCO beads were synthesized in house: NHS-activated Sepharose™ fast flow (#17-0906-01, GE Healthcare) was incubated to varying concentrations of dibenzocyclooctyne-amine (DBCO-amine, #761540, Sigma-Aldrich). The prepared beads were stored as 50% slurry in a 1:1 ethanol: water mixture. AF488-Azide (#CLK-1275-1, Jena Bioscience) was used to test success of bead – DBCO coupling. Trypsin gold was purchased from Promega (Mannheim, Germany) and lysyl endopeptidase (LysC) was from Wako (Neuss, Germany). Benzonase® - pharmaceutical production purity - was from Merck (Darmstadt, Germany).

### Procedure

A schematic overview of the workflow is shown in Figure 1 and described in detail below:

**Figure 1:**
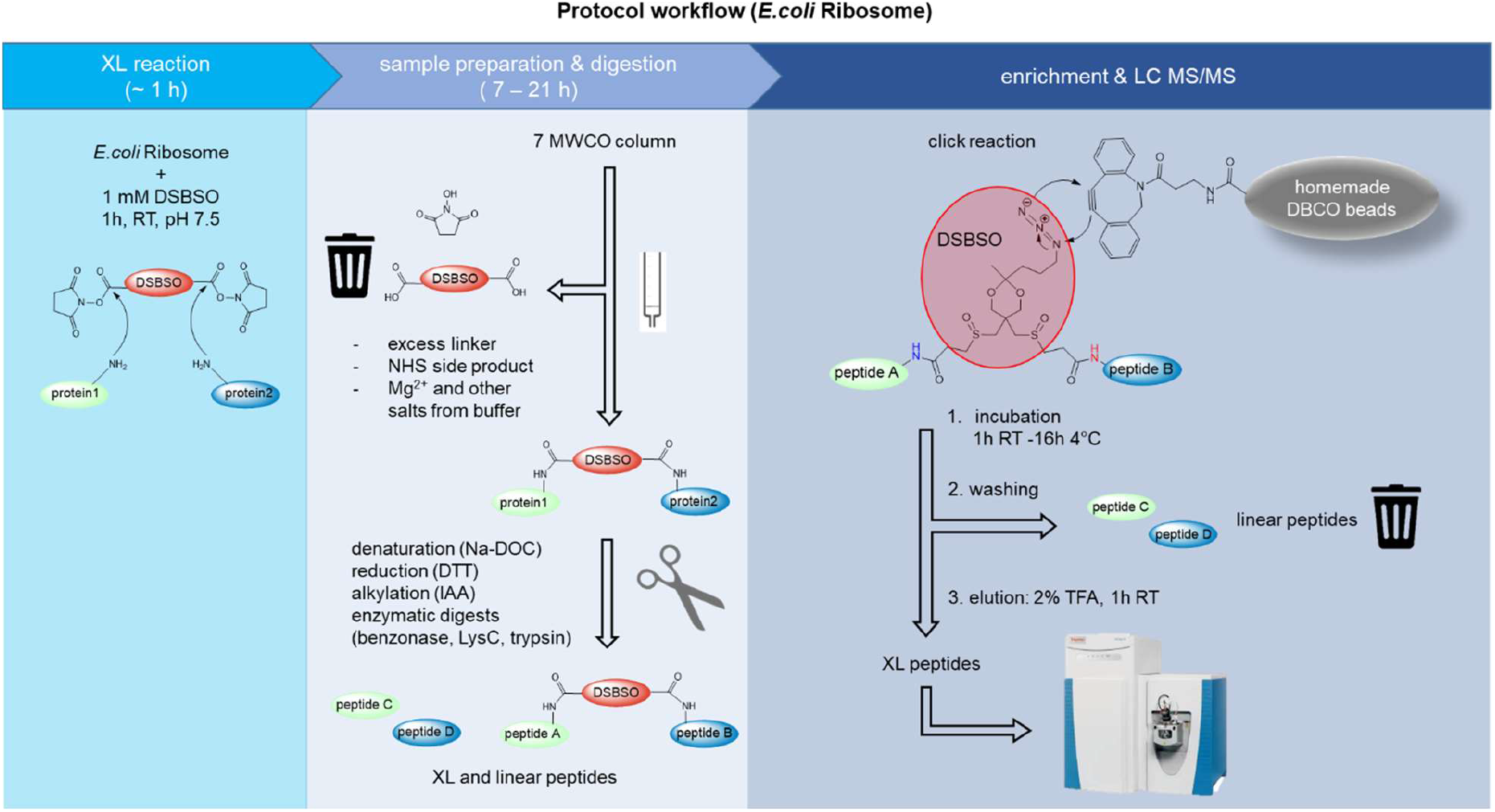
Schematic workflow of DSBSO enrichment method using DBCO coupled Sepharose beads on the example of *E. coli* ribosome.

#### XL reaction

*E. coli* ribosome was diluted with dilution buffer to a final concentration of 1 mg/mL. DSBSO linker was dissolved in dry DMSO to produce a 40x stock solution immediately prior to use. For *E. coli* ribosome a 40 mM stock solution was prepared. The cross-linking reaction was initiated by addition of DSBSO stock solution in a final concentration of 1 mM to the diluted *E. coli* ribosome and incubated for 1 h at 25 °C with mild agitation. Quenching was performed by addition of 1 M Tris pH 7.5 reaching a final concentration of 100 mM Tris. Incubate for 15 min at 25 °C with mild agitation.

Removal of excess linker by exchanging the buffer to 50 mM HEPES pH 7.5 was done by use of a Zeba Spin Column, according to manufacturer’s recommendation.

#### Protein preparation and digestion

Denaturation was induced by addition of a 15% sodium deoxycholate (Na-DOC) solution reaching a final concentration of 1.5% Na-DOC. Reduction was performed using DTT at a final concentration of 10 mM. Additionally, 0.5 μL benzonase was added to degrade nucleotides and the mixture was incubated for 60 min at 37 °C. Alternatively, if no benzonase digestion was needed (Cas9 samples), the mixture was incubated for 30 min at 56 °C. Alkylation was performed using IAA (iodacetamide) at a final concentration of 20 mM and the mixture was incubated for 30 min at room temperature in the dark. Quenching was performed by addition DTT (dithiothreitol) at a final concentration of 5 mM and incubation for 15 min at room temperature. For sequential digestion the sample mixture was diluted using 50 mM HEPES pH 7.5 to reach a concentration of 1 % Na-DOC. Subsequently Lysyl Endopeptidase (LysC) in a 1:50 (w/w) ratio was added and incubated for 2 h at 37 °C. Subsequently trypsin, in a 1:50 (w/w) ratio, was added and incubated for further 16 h at 37 °C. For quality control purposes, we checked the success of the digestion by injection of a small sample aliquot to an HPLC system and compare it to a fully digested reference.

#### Enrichment of XL -peptides

To compare the performance of the DBCO beads to an established enrichment method for DSBSO linked peptides, enrichment was alternatively also performed similar, as described by Kaake *et al.*^15^ with the following details: The denatured protein was incubated to 100 μM BARAC biotin over night at 4 °C. Excess BARAC biotin was removed by use of the Zeba Spin Column as before. Further sample preparation was performed as for the DBCO method (reduction, alkylation, digestion). Enrichment was performed by incubation to streptavidin beads, which were subsequently washed as described for the DBCO method and eluted using an aqueous mixture of 20% formic acid and 10% acetonitrile as described by Kaake et al.^15^ This method is indicated as “BARAC biotin method” within this publication.

For the, here established, DBCO method the optimal excess ratio of DBCO groups (immobilized on beads) to azide groups (based in used input of DSBSO cross-linker) was estimated to be 10x, which corresponded to 12 μL bead slurry.

For experiments with HEK peptide background added, tryptically digested HEK peptides were added prior to incubation to enrichment beads (DBCO beads, or streptavidin) in the given ratios.

The DBCO beads were equilibrated by washing them 3x using at least 5 bead volumes of 50 mM HEPES pH 7.5 buffer. The beads were separated by centrifugation at 2000 g for 1 min each and the supernatant was carefully removed without disruption of the beads. The prepared XL sample was mixed with equilibrated DBCO-beads and allowed to react for at least 1 h at 25 °C with gentle agitation. Alternatively, incubation was performed over night at 4°C without affecting enrichment performance. The remaining supernatant was removed and stored to check for successful click reaction. The beads were washed using at least 5 bead volumes as follows and the beads were separated from the washing solution after each step by centrifugation at 200 g for 1 min: 3x washing with 50 mM HEPES pH 7.5, 1 M NaCl, 3x washing with 10% ACN in H_2_O and finally 3x washing with 10 mM Tris pH 7.5. Elution of XL-peptides from the beads was performed by acidic cleavage of the acetal bond within DSBSO using the same volume of 2% (v/v) trifluoracetic acid (TFA) in H_2_O as used as input bead-slurry volume. After incubation for at least 1 h at 25°C, the eluate was separated from the beads and transferred into fresh tubes containing 5 % (v/v) of DMSO based on the final volume. (Addition of DMSO is optional, however we have noticed that XL-peptides tend to stick to the walls of tubes, and we were therefore able to increase the number of detectable XLs by addition of DMSO as a solvent)

#### Mass spectrometry

After acidic elution from the beads, the samples were subjected to LC ‒MS/MS analysis without any freeze/thaw cycle in between. Control samples, where no enrichment strategy was applied, were prepared and digested as explained above but not incubated to any beads, followed by acidification using 10% (v/v) TFA finally reaching 1% (v/v) TFA.

Enriched- and control-samples were separated using a Dionex UltiMate 3000 HPLC RSLC nano-system coupled to an Q Exactive™ HF-X Orbitrap mass spectrometer via Proxeon nanospray source or to an Orbitrap Fusion™ Lumos™ Tribrid™ mass spectrometer EASY ESI source (all: Thermo Fisher Scientific). Samples were loaded onto a trap column (Thermo Fisher Scientific, PepMap C18, 5 mm × 300 μm ID, 5 μm particles, 100 Å pore size) at a flow rate of 25 μL min^−1^ using 0.1 % TFA as mobile phase. After 10 min, the trap column was switched in line with the analytical column (Thermo Fisher Scientific, PepMap C18, 500 mm × 75 μm ID, 2 μm, 100 Å). Peptides were eluted using a flow rate of 230 nl min^−1^, with the following gradient over 80 min or 110 min for Cas9 and *E. coli* ribosome samples respectively: 0 −10 min 2 % buffer B, followed by an increasing concentration of buffer B up to 35 % or 40 % until min 60 or 90 for Cas9 or *E. coli* ribosome samples respectively. This is followed by a 5 min gradient from reaching 95 % B, washing for 5 min with 95% B, followed by re-equilibration of the column at 30°C (buffer B: 80 % ACN, 19.92 % H_2_O and 0.08 % TFA, buffer A: 99.9% H_2_O, 0.1% TFA).

The mass spectrometer was operated in a data-dependent mode, using a full scan (m/z range 375-1500, nominal resolution of 120.000, target value 1E6). MS/MS spectra were acquired by stepped HCD using an NCE (normalized collision energy) of 27±6, an isolation width of 0.8 m/z, a resolution of 30.000 and the target value was set to 5E4. Precursor ions selected for fragmentation (± 10 ppm, including exclusively charge states 3-8) were put on a dynamic exclusion list for 20 s. Additionally, the minimum AGC target was set to 5E4 and precursors with highest charges were given priority. Measurements on the Q Exactive™ HF-X Orbitrap were performed with similar settings and following details changed: m/z 350-1600, isolation width 1 m/z, intensity threshold 3.3E4/AGC 5E3.

#### Data analysis

MS data were analyzed with the help of Thermo Proteome Discoverer (2.3.0.523). Peptide identification was performed by MS Amanda (2.3.0.12368)^26^. The peptide mass tolerance was set to ±5 ppm and the fragment mass tolerance to ±0.02 Da. Carbamidomethyl (+57.021 Da) at cysteine was set as static modification. Oxidation (+15.995 Da) at methionine was set as dynamic modification. The result was filtered to 1% FDR (false discovery rate) on peptide level using the Target Decoy PSM Validator integrated in Thermo Proteome Discoverer, PSM hits were additionally filtered for a minimum score of 150.

Cross-links were identified either using XlinkX 2.2^27^ as node within Proteome Discoverer (2.3.0.523) or using MeroX (2.0.0.6)^7^ as indicated. For XlinkX the cross-link modification DSBSO was defined as C_11_H_16_O_6_S_2_ with the following crosslink-fragments: Alkene C_3_H_2_O, thiol C_8_H1_2_O_4_S_2_, sulfenic acid C_8_H_14_O_5_S_2_ and linker specificity towards lysine and N-terminal amino residues was set. Fixed carbamidomethylation of cysteine and variable oxidation of methionine residues were set as modifications. Standard settings were used, with a minimal XlinkX score of 40 and a minimal delta score of 4 and results were filtered at 5 % FDR at peptide level using the XlinkX validator node. To analyze data with MeroX, raw files were first converted to mgf format using MSConvertGUI (3.0.19085-a306312d7)^28^ without using any filter. Data analysis was performed using the following settings: C-terminal cleavage sites lysine and arginine with 3 missed cleavages, allowed peptide length: 5 to 30, as static modification acetamidation of cysteine and as variable modification oxidation of methionine was set. The cross-linker DSBSO was defined as follows: DSBSO with specificity towards lysine and N-termini, fragments at site 1 and 2: Alkene and Thiol, as given for XlinkX above. Additionally, the following diagnostic ions were set: C_8_H_12_NO, C_8_H_15_N_2_O, C_8_H_12_NOS, C_8_H_15_N_2_OS. Dead ends were allowed to react with H_2_O. Further settings: precursor precision 4 ppm, fragment ion precision 8 ppm, S/N ratio 1.5, precursor masses were corrected (max 3 isotope shifts). Prescore intensity 10 %, FDR cutoff 5 %, score cutoff −1, for analysis with large databases, the proteome wide mode with a minimum peptide score of 10 was used, otherwise RISEUP mode was used.

Quantification of cross linked peptides was performed label free using apQuant (3.1.1.27312)^29^ within Proteome Discoverer 2.3.

As database for Cas9 samples the sequence of S. Pyogenes Cas9 with fused HaloTag® plus the full human Swiss-Prot (as of 13^th^ March 2018, 20271 proteins) was used. For ribosome samples a shotgun database containing 171 proteins, generated earlier^9^, was used and in experiments spiked with tryptic HEK peptides the full human Swiss-Prot (as of 13^th^ March 2018, 20271 proteins) was added.

The mass spectrometry proteomics data have been deposited to the ProteomeXchange Consortium via the PRIDE^30^ partner repository with the dataset identifier PXD016963

## Results and Discussion

### Generation and evaluation of DBCO coupled Sepharose beads

We generated DBCO coupled beads by reaction of DBCO to NHS preactivated Sepharose™. To test their loading capacity, we performed a click reaction to an Alexa488 tagged azide. By photometric fluorescence detection, we estimated a loading of ~ 5.9 μmol DBCO groups/ mL Sepharose-bead-slurry.

Next, we evaluated cleavage conditions for the labile acetal functionality on the linker. TFA at a concentration of 2 % (v/v) for 1 h at 25 °C, was thereby sufficient to cleave the azide tag off from DSBSO. This hydrolysis step thereby reached close to 100 % yield (estimated by MS on a LTQ Orbitrap Velos). Importantly, these cleavage conditions are milder compared to overnight incubation with 20 % formic acid, 20 % acetonitrile as originally used by Kaake et al.^15^ To probe MS compatibility of the synthesized DBCO-beads, empty beads were incubated to 2 % TFA for 1 h without producing any appreciable background signal within MS. In contrast, commercially available DBCO coupled beads generated interfering background signals.

To test for the optimal amount of bead material to be used, 20 μg Cas9 protein were cross-linked with 0.5 mM DSBSO, reduced, alkylated, digested with trypsin and incubated to varying amounts of DBCO beads (Figure 2A). All data were evaluated using XlinkX^27^ and MeroX^7^ to increase confidence in the numbers of cross-links reported throughout this study and those data were analyzed against the human Swiss-Prot supplemented with Cas9 (see experimental section) to be comparable to the spike-experiments in Figure 2C. Most unique cross-links can be detected when using 12 μL of DBCO beads (Figure 2A), which corresponds to a 10x excess ratio of DBCO groups (on the beads) to azide (corresponding to the amount of DSBSO linker added to the sample). The co-enriched mono-linked peptides show a comparable pattern as seen for cross-link numbers (Figure 1B). When using 24 μL bead material, unspecific peptide binding or less effective elution from the huge bead excess, however, led again to slightly lowered cross-link numbers. A slightly increased peptide background, in case of large bead amounts used, is indeed visible (Figure 2B). While we detected up to 465 (XlinkX) unique cross-links on Cas9 after enrichment (originating from 20 μg linked Cas9), in a control only 96 links were detectable (Figure 2A). Of note, a maximum amount of 1 μg total protein was injected for each control run to prevent column overloading. In contrast, 100 % of enriched samples, originating from 20 μg input, could be loaded without any risk of overloading. Through enrichment, both selectivity and sensitivity were improved. To estimate an enrichment factor, we therefore formed the ratio of cross-linked peptides (cross-link sequence matches via XlinkX) over linear peptides (peptide sequence matches, excluding DSBSO modified peptides, via MS Amanda). This factor is clearly increased after enrichment, compared to the control and the resulting bars show a similar trend as the unique cross-link numbers (Supplemental Figure 1). In an additional experiment we were interested in uncaptured cross-linked peptides remaining in the depleted fraction after enrichment. We analyzed the depleted fraction after 1 - 3 repeated incubations to fresh DBCO beads. In parallel the beads obtained from these 1 −3 incubations were pooled respectively and the enriched cross-linked peptides were analyzed to see if additional links can be detected after multiple incubation to the beads (Supplemental Figure 2). In this experiment the number of unique cross-linked peptides and mono-linked peptides was not increased upon multiple incubations, although both numbers were decreased upon multiple incubation to beads in the remaining depleted fraction (Supplemental Figure 2 A and B). As shown in Supplemental Figure 2 C the overlap of detected unique cross-links within the enriched and depleted fraction after a single enrichment step is very high. This demonstrates that a single incubation to DBCO beads is already efficient to enrich all detectable cross-links.

**Figure 2:**
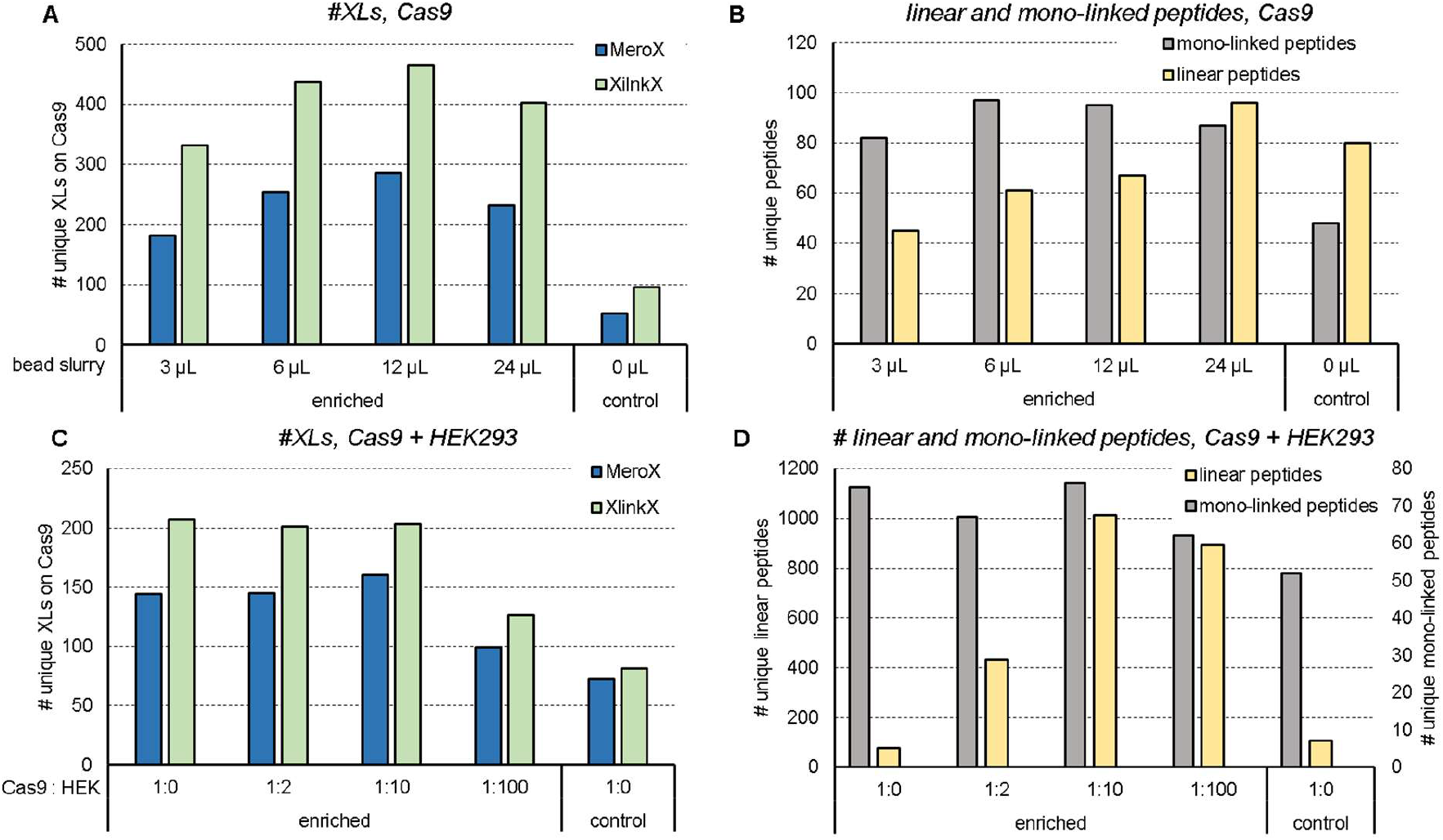
Optimization of input bead amount and probing of the enrichment method based on Halo-tagged recombinant Cas9 protein. Number of unique cross-links within Cas9 (**A**) or unique mono-linked peptides (as given by MeroX) and linear peptides (as given by MS Amanda) (**B**) after linking 20 μg of the recombinant protein with 0.5 mM DSBSO with or without enrichment (control) using the indicated bead slurry volumes. Number of unique cross-links on Cas9 (**C**), or unique detected mono-linked and linear peptides (**D**) after linking 20 μg Cas9 each and spiking into tryptic HEK peptides in excess as indicated, enriched using 12 μL of DBCO bead slurry each. Results filtered for 5 % FDR, n=1

Overall, in our study, the ideal bead-slurry volume was estimated to be 6 −12 μL, corresponding to a 5x to 10x excess of DBCO groups over azide groups respectively and a single incubation to DBCO beads is enough to enable a sufficient enrichment with a very high coverage of present cross-linked peptides. Based on these results 12 μL bead slurry / 20 μg cross-linked input material was used for all further experiments.

### Enrichment of Cas9 in a complex environment

In a next step, we spiked cross-linked Cas9 peptides into a background of tryptic HEK peptides (Figure 2C, D). Previous experiments already showed that no cross-links were detectable after spiking 1:2 (20 μg Cas9 + 40 μg HEK) without using an enrichment strategy. In contrast, when using our enrichment method, the number of detectable links upon diluting the sample with an increasing HEK background remains close to its original number (without HEK peptide addition, 1:0, Figure 2C). Figure 1D shows, that the background of linear peptides, which are mainly originating from the added HEK peptides, increases with increasing spike-ratios. This indicates some unspecific binding to the beads, which could not be washed away, however, the numbers do not further increase from 1:10 to 1:100 spike ratio although 10x more background was initially added.

### Probing the enrichment method in a more complex system – *E. coli* ribosome

Aiming to increase the complexity of our model system we decided to use *E. coli* ribosome (NEB, MA, USA) which was finally also spiked into a tryptic HEK peptide background. For those experiments ribosomal proteins were linked using 1 mM DSBSO for 1 h. After reduction, alkylation and digestion, the obtained cross-linked peptides were enriched by incubation with 10 eq excess of DBCO beads either for 1 h at 25 °C or overnight at 4°C. Elution was performed by incubation to 2% TFA for 1 h, prior to measurement via LC-MS. (A schematic workflow shown in the graphical abstract.) Especially in the case of more complex samples we observed, that separation of hydrolyzed linker and side products of the cross-linking reaction by use of a Zeba Spin 7 MWCO column yielded slightly improved final unique cross-link numbers (while it did not change our results for simple proteins like Cas9). This is most likely due to less free DSBSO ‒not covalently bound to any protein – and thus not consuming DBCO groups on the beads. For *E. coli* ribosome, recovery of protein after elution from the Zeba Spin column was estimated to be ~ 95 % based on detection at 214 nm after HPLC separation. This additional step furthermore separates Mg^2+^ ions, therefore hindering sodium-deoxycholate from forming a precipitate, which would impair denaturation. Denaturation of ribosomal protein samples was performed to improve digestion efficiency. We also tested to denature with urea or omit denaturation. This, independently of using the Zeba Spin column, led to lower cross-link numbers compared to using deoxycholate.

The number of unique cross-links detected within the ribosome was more than doubled from 47 before-to 109 links after-enrichment (Figure 3A) on average (XlinkX). An additional comparison of this data analyzed with a 1 % FDR instead of 5 % FDR cutoff is shown in Supplemental Figure 3. The picture seen on cross-link numbers, is mirrored when looking on mono-linked peptides. In line with this data, the number of unique linear peptides was strongly decreased in the enriched fraction (Figure 3B). Furthermore, we again estimated the enrichment factor (as described for Cas9 above), which indicated an effective enrichment (Figure 3C).

**Figure 3:**
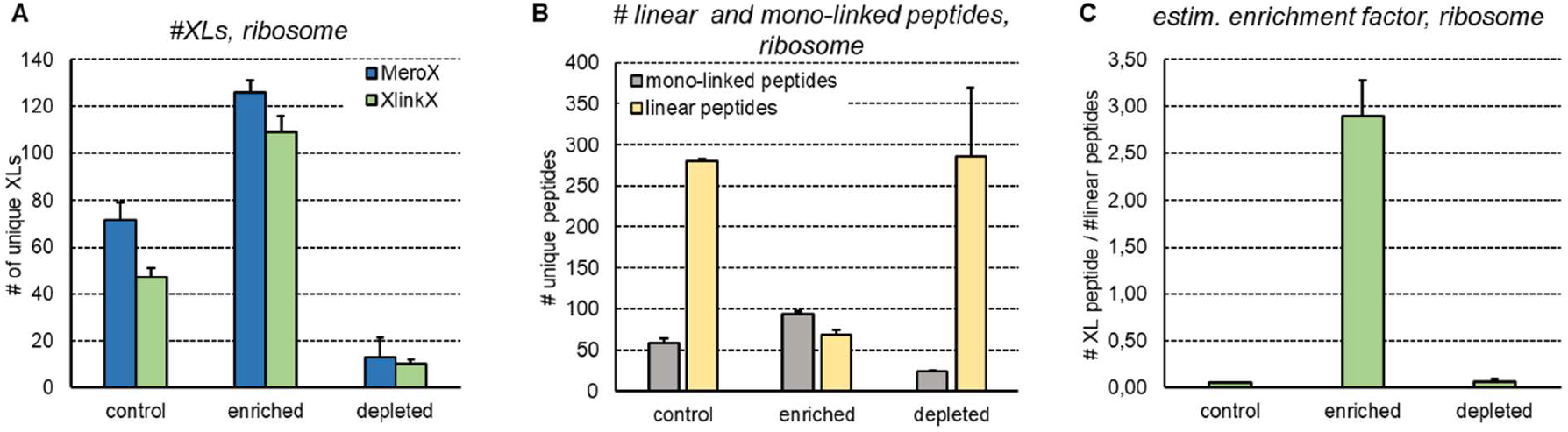
Probing enrichment on a more complex system using cross-linked *E. coli* ribosome. Number of unique cross-links detected within ribosome-shotgun database (**A**), number of detected unique linear peptides without a DSBSO modification from MS Amanda and number of unique mono-linked peptides from MeroX (**B**), ratio of detected cross-linked (XlinkX) over linear peptides (MS Amanda) (**C**) without enrichment (control), after DBCO bead enrichment (enriched) or in the remaining supernatant over the beads after click reaction (depleted). 20 μg *E. coli* ribosome each were linked using 1 mM DSBSO. Bars indicate the average values, with standard deviation depicted as error bar, 5 % FDR, n≥3.

Of note, the depleted fraction still contained some cross-links. As already seen in case of Cas9 (Supplemental Figure 2 C), the majority of those cross-links was also detected in the enriched fraction (Supplemental Figure 4 A, Supplementary Table 1). We analyzed the relative ion intensity within the enriched fractions of those cross-links also found in the depleted fraction, showing that predominantly high abundant cross-links were also found in the depleted fraction. Within the enriched fraction those cross-links make up ~40 % of the total ion intensity (but only ~ 9 % of the total number). In total the ion intensity of all detected cross-links is only 1/1200 in the depleted vs control fraction (1 μg injected each, see Supplemental Figure 4 B). In contrast, the relative ion intensity of detected proteins remained on the same level (Supplemental Figure 4 C) showing that they were not unspecifically captured. The performance of the enrichment method becomes most obvious when doing the same experiment within a background of tryptic HEK peptides, with no cross-link ions detectable at all in the depleted fraction but a similar ion intensity as without spiking in the enriched fraction (Supplemental Figure 4 D). The amount of non-cross-linked material was thereby again clearly reduced by enrichment (Supplemental Figure 4 E). The generally observed increase in cross-link numbers upon enrichment is mainly reasoned by a reduction of background, whose intensity was reduced by factor 611 although 20x more input was used (corresponds to a reduction by factor 12000, see Supplemental Figure 4 C). In conclusion, the possible input amount can be significantly increased without overloading the column, to boost cross-link numbers: In detail, the relative ion intensity of cross-linked peptides vs total intensity was on average 0.47 % prior to enrichment., while merely any linear peptides remained after enrichment, yielding to an average ion intensity of 93.2 % cross-linked peptides after enrichment (Supplemental Figure 5A). As for the experiments with Cas9, for control samples 1 μg of total protein was injected each, while we were able to load 100% of each eluate after enrichment. Since we used 20 μg input material for all enrichment experiments, this would correspond to roughly 100 ng of cross-linked peptides (based on 0.5% estimated abundance of XL peptides). Since this is still a relatively tiny injection amount, we increased the input for enrichment from 20 to 100 μg linked ribosomal proteins in a single experiment. This led to a TIC (total ion current) increase by factor 3 to ~ 6E9 and we detected 215 unique cross links via XlinkX (increased by factor 2, Supplemental Figure 5B). The relative ion intensity of cross-linked material before and after enrichment thereby remained on a similar level as for lower inputs (Supplemental Figure 5C). In conclusion, due to the relatively high purity of enriched cross-linked peptide samples, cross-link numbers could likely be increased for all experiments in this study if higher sample amounts would have been used.

### Recovery check on cross-linked *E. coli* ribosome

Additionally, we decided to probe our method by mimicking an *in vivo* system and spiked linked ribosome into a tryptic HEK digest in up to 100-fold excess. No links can be detected any more after spiking, if the excess of HEK peptides is exceeding 2-fold. In contrast, we were still able to recover 106 (MeroX) / 73 (XlinkX) unique XLs based on 5 % FDR when spiking 1:100 into HEK peptides. (Figure 4 A, B, Supplemental Figure 4 D). Compared to the BARAC-biotin approach we were able to get ~5 x increased cross-link numbers (unspiked condition). The recovery of cross-links was also significantly improved by means of LFQ: The relative ion intensity of cross-linked peptides after enrichment of (unspiked) samples was > 95 %, compared to ~ 58% when using the two-step BARAC-biotin method. Additionally, mono-linked peptides were analyzed by MeroX, showing that they are also enriched by using the DBCO method (Figure 4C). The background of linear peptides, however, was also increased with increasing spiking ratios and reached the same level at 1:100 as for controls with 1:2 spike ratio (data see Figure 4D; As mentioned before, the enriched samples originate from 20x more input material. Therefore, similar peptide levels still indicate a significant reduction in linear peptide concentration.). Especially for highly diluted samples more and longer washing steps might still decrease that background. In our hands, however, this did not significantly boost the obtained cross-link numbers. As alternative to washing with 1 M NaCl in HEPES buffer and 10 % ACN in water, we also tried to reduce background signal by washing with up to 2 % SDS, 4 M urea or 4 M guanidinium hydrochloride. This led to slightly reduced background signals but the number of detected final cross-links was again not increased. Another reason for slightly decreased cross-link numbers upon high spike ratios, might be an incomplete click reaction due to a highly diluted linker.

**Figure 4:**
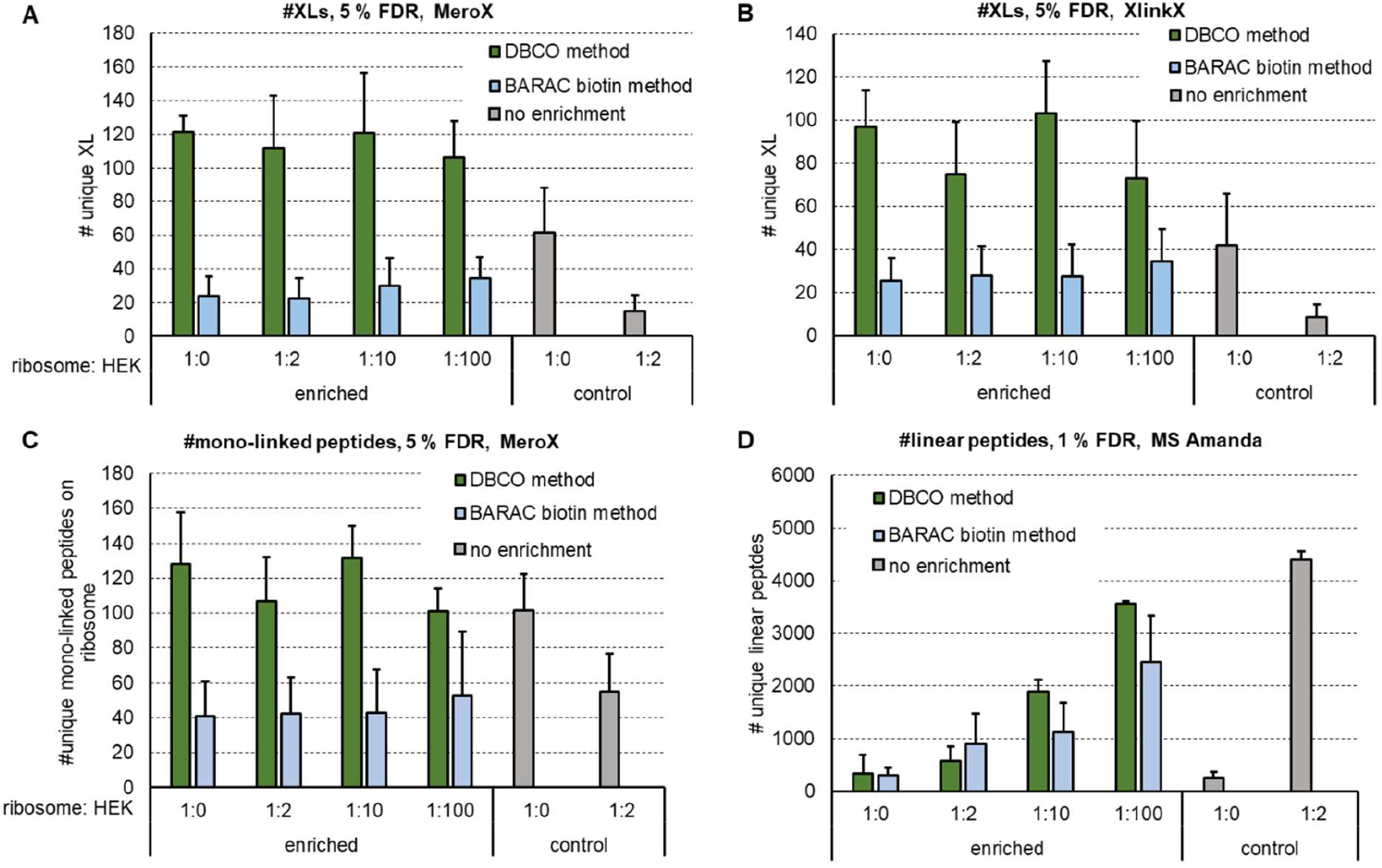
Check for recovery of DBCO enrichment compared to the published BARAC biotin method by increasing a tryptic HEK background to mimic a cellular environment. Number of detected unique cross-links using MeroX (**A**) or XlinkX (**B**) for analysis, number of detected unique mono-linked peptides via MeroX (**C**) and unique linear peptides via MS Amanda (**D**) after cross-linking 20 μg of *E. coli ribosome* each using 1 mM DSBSO. Tryptic HEK peptides were added to each sample in the given excess prior to enrichment with DBCO beads. Bars indicate the average values, with standard deviation depicted as error bar, FDR as indicated, n≥3.

Finally we estimated the correctness of the software calculated FDR rate of the obtained data, by separating cross-link hits found within the ribosomal shotgun proteins^9^ and found within the human proteome from our searches using the combined database. As shown in Supplemental Figure 6, for XlinkX, the number of “wrong” cross-links from or to human proteins varies between 4 – 8 % of the total hit number and is therefore in the expected range. The actual FDR, however, might still be higher, reasoned by potential false positive cross-links within the ribosome.

We are aware that – by means of obtained absolute XL numbers – others have reported more unique cross-links (up to 766 unique links for *E.coli* ribosome using DSSO or DSBU and fractionation using SEC or SCX respectively^9,31^). Enriching via an affinity tag may still be very advantageous in case of a complex matrix, as this would be the case for *in vivo* investigations. The ability of the DBCO method to work in a cell lysate or potentially also *in vivo* was successfully investigated by spiking cross-linked material into HEK peptides and DSBSO was already shown to be cell permeable by the inventors^15^ of the cross-linker.

### Sensitivity check on cross-linked *E. coli* ribosome

In addition to looking at the recovery of XLs we checked for the sensitivity of the enrichment method, by spiking decreasing amounts of cross-linked ribosomal peptides into a constant background of 100 μg tryptic HEK peptides (Figure 5). In Figure 5, the given ratios are calculated based on the original protein amount used for XL reaction, meaning e.g. for 0.25:100, 0.25 μg *E. coli* ribosomal protein were used for DSBSO linking and prior to incubation with the beads for click-reaction 100 μg tryptic HEK peptides were added. Our sensitivity data shows, that the here described method is indeed very sensitive, due to the high selectivity of the click reaction. Cross-linked peptides are pulled from a mixture with extreme low concentrations of cross-linked material: As mentioned above, we estimate that ~0.47 % of the ribosomal peptides are cross-linked prior to enrichment, leading to the assumption, that an ~ 1 ng of cross-linked peptides (based on 0.47 % estimated abundance of successfully linked peptides and 250 ng of protein input) in 100 μg of linear peptides was still sufficient for detection of some unique cross-links (Figure 5A). Furthermore, again, mono-links are co-enriched. The visible background of linear peptides is reduced after enrichment and remains constant for all conditions, which is a consequence of equal amounts of tryptic HEK peptides added to each sample (Figure 5B).

**Figure 5:**
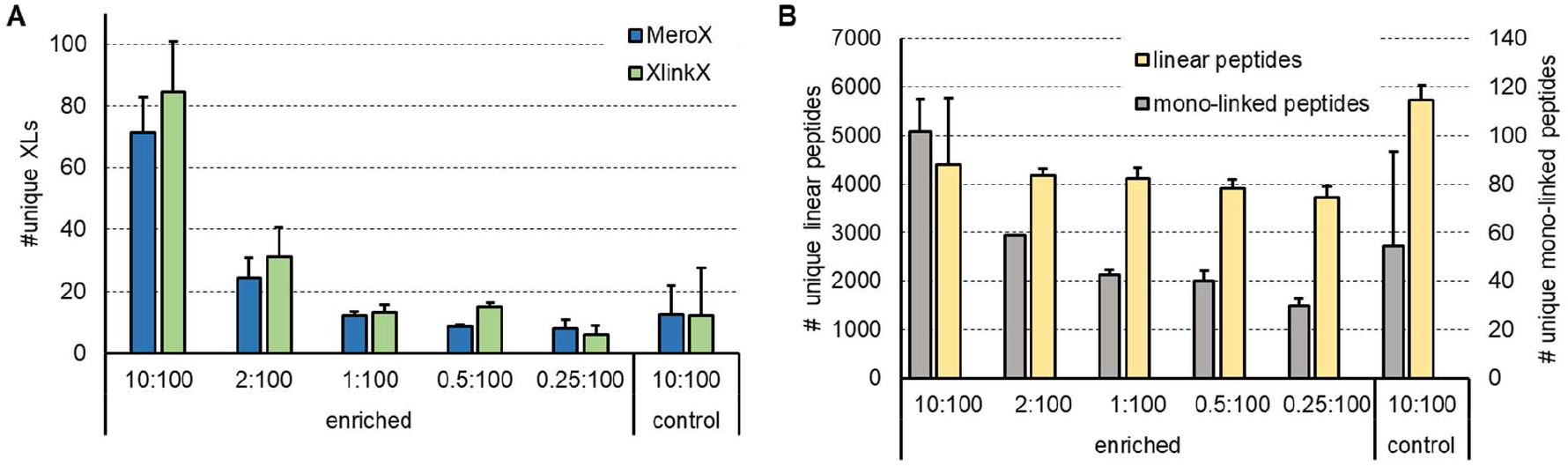
Check for sensitivity of DBCO enrichment by decreasing the amount of XL material within a constant tryptic HEK background to mimic a cellular environment. Number of detected unique cross-links via MeroX and XlinkX respectively (**A**) and number of detected unique mono-linked (MeroX) and unique linear peptides (MS Amanda) (**B**), after cross-linking *E. coli* ribosome using 1 mM DSBSO. 100 μg tryptic HEK peptides were added to 0.25 −10 μg linked ribosome as indicated prior to enrichment with DBCO beads. Bars indicate the average values, with standard deviation depicted as error bar, 5 % FDR, n = 2.

## Conclusion

The introduced DBCO bead affinity enrichment provides an effective and simplified method for purification of cross-linked peptides utilizing a biorthogonal and therefore selective click chemistry reaction.

Although further optimization and improvements must be done in future experiments, we believe, that the published protocol will be of great value to other groups aiming to enrich cross-linked samples. This will especially be an advantage for larger protein complexes and when linking *in vivo* or on beads during an immuno-precipitation. The synthesis of the beads is thereby easy and quick. Additionally, this one step method omits the use of other, time consuming, fractionation and enrichment steps. In theory the method can also be used for a simplified enrichment of other azide-tagged cross-linkers (e.g. cliXlink^32^), analysis tools (e.g. DYn-2 for analysis of protein-S-sulfenylation^33^) or biomolecules (e.g. incorporated azidonucleosides^34^) and will therefore be of great value for in vitro and *in vivo* studies using proteins or DNA with the respective mutations or modifications bearing a bio-orthogonal azide tag.

## Acknowledgement

The authors thank Kristina M Uzunova (molecular biology service group, IMP/IMBA/GMI) who prepared the DBCO coupled Sepharose beads for us. Furthermore, our gratitude goes to Elisabeth Roitinger for her continuous support and fruitful discussions as well as to Rebecca Beveridge and Johannes Stadlmann for proofreading the manuscript.

## Supporting Information

The following supporting information is available free of charge at the ACS website http://pubs.acs.org

Figure S1: Estimated enrichment factor. Figure S2: Multiple incubation of DSBSO linked Cas9 to DBCO beads. Figure S3: Enrichment of *E. coli* ribosomal cross-linked peptides. Figure S4: A closer look to the enrichment using cross-linked *E. coli* ribosome. Figure S5: Higher protein input is possible after enrichment, leading to more cross-links. Figure S6: Error estimation – spiked *E. coli* ribosome samples.

## For TOC only

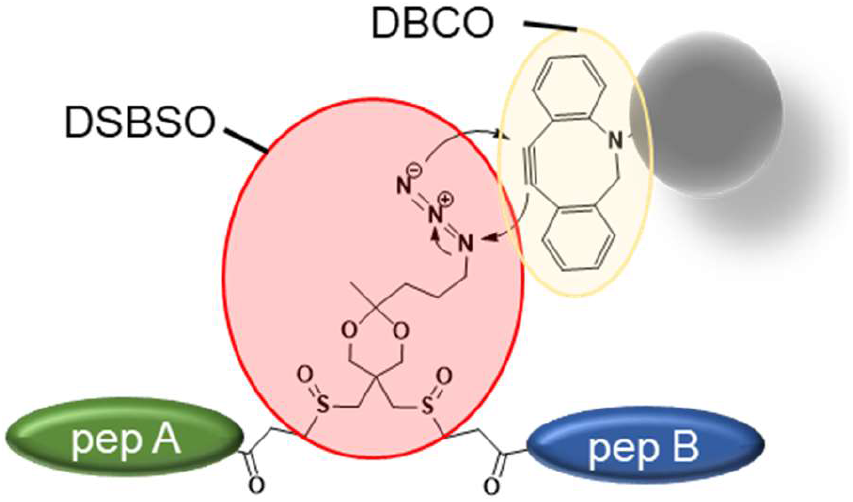

## References

(1) Leitner, A.; Faini, M.; Stengel, F.; Aebersold, R. Crosslinking and Mass Spectrometry: An Integrated Technology to Understand the Structure and Function of Molecular Machines. Trends in Biochemical Sciences 2016, 41 (1), 20–32. https://doi.org/10.1016/j.tibs.2015.10.008.

(2) Rappsilber, J. The Beginning of a Beautiful Friendship: Cross-Linking/Mass Spectrometry and Modelling of Proteins and Multi-Protein Complexes. J. Struct. Biol. 2011, 173 (3), 530–540. https://doi.org/10.1016/j.jsb.2010.10.014.

(3) Yu, C.; Huang, L. Cross-Linking Mass Spectrometry (XL-MS): An Emerging Technology for Interactomics and Structural Biology. Anal Chem 2018, 90 (1), 144–165. https://doi.org/10.1021/acs.analchem.7b04431.

(4) Chavez, J. D.; Bruce, J. E. Chemical Cross-Linking with Mass Spectrometry: A Tool for Systems Structural Biology. Current Opinion in Chemical Biology 2019, 48, 8–18. https://doi.org/10.1016/j.cbpa.2018.08.006.

(5) Leitner, A.; Walzthoeni, T.; Kahraman, A.; Herzog, F.; Rinner, O.; Beck, M.; Aebersold, R. Probing Native Protein Structures by Chemical Cross-Linking, Mass Spectrometry, and Bioinformatics. Mol Cell Proteomics 2010, 9 (8), 1634–1649. https://doi.org/10.1074/mcp.R000001-MCP201.

(6) Steigenberger, B.; Pieters, R. J.; Heck, A. J. R.; Scheltema, R. A. PhoX: An IMAC-Enrichable Cross-Linking Reagent. ACS Cent. Sci. 2019, 5 (9), 1514–1522. https://doi.org/10.1021/acscentsci.9b00416.

(7) Götze, M.; Iacobucci, C.; Ihling, C. H.; Sinz, A. A Simple Cross-Linking/Mass Spectrometry Workflow for Studying System-Wide Protein Interactions. Anal. Chem. 2019, 91 (15), 10236–10244. https://doi.org/10.1021/acs.analchem.9b02372.

(8) Mendes, M. L.; Fischer, L.; Chen, Z. A.; Barbon, M.; O’Reilly, F. J.; Giese, S. H.; Bohlke-Schneider, M.; Belsom, A.; Dau, T.; Combe, C. W.; et al. An Integrated Workflow for Crosslinking Mass Spectrometry. Mol. Syst. Biol. 2019, 15 (9), e8994. https://doi.org/10.15252/msb.20198994.

(9) Stieger, C. E.; Doppler, P.; Mechtler, K. Optimized Fragmentation Improves the Identification of Peptides Cross-Linked by MS-Cleavable Reagents. J. Proteome Res. 2019, 18 (3), 1363–1370. https://doi.org/10.1021/acs.jproteome.8b00947.

(10) Leitner, A.; Reischl, R.; Walzthoeni, T.; Herzog, F.; Bohn, S.; Förster, F.; Aebersold, R. Expanding the Chemical Cross-Linking Toolbox by the Use of Multiple Proteases and Enrichment by Size Exclusion Chromatography. Molecular & Cellular Proteomics 2012, 11 (3), M111.014126. https://doi.org/10.1074/mcp.M111.014126.

(11) Klykov, O.; Steigenberger, B.; Pektaş, S.; Fasci, D.; Heck, A. J. R.; Scheltema, R. A. Efficient and Robust Proteome-Wide Approaches for Cross-Linking Mass Spectrometry. Nat Protoc 2018, 13 (12), 2964–2990. https://doi.org/10.1038/s41596-018-0074-x.

(12) Tinnefeld, V.; Venne, A. S.; Sickmann, A.; Zahedi, R. P. Enrichment of Cross-Linked Peptides Using Charge-Based Fractional Diagonal Chromatography (ChaFRADIC). J. Proteome Res. 2017, 16 (2), 459–469. https://doi.org/10.1021/acs.jproteome.6b00587.

(13) Fritzsche, R.; Ihling, C. H.; Götze, M.; Sinz, A. Optimizing the Enrichment of Cross-Linked Products for Mass Spectrometric Protein Analysis. Rapid Communications in Mass Spectrometry 2012, 26 (6), 653–658. https://doi.org/10.1002/rcm.6150.

(14) Fürsch, J.; Kammer, K.-M.; Kreft, S. G.; Beck, M.; Stengel, F. Proteome-Wide Structural Probing of Low-Abundant Protein Interactions by Crosslinking. bioRxiv 2019, 867952. https://doi.org/10.1101/867952.

(15) Kaake, R. M.; Wang, X.; Burke, A.; Yu, C.; Kandur, W.; Yang, Y.; Novtisky, E. J.; Second, T.; Duan, J.; Kao, A.; et al. A New in Vivo Cross-Linking Mass Spectrometry Platform to Define Protein– Protein Interactions in Living Cells. Molecular & Cellular Proteomics 2014, 13 (12), 3533–3543. https://doi.org/10.1074/mcp.M114.042630.

(16) Weisbrod, C. R.; Chavez, J. D.; Eng, J. K.; Yang, L.; Zheng, C.; Bruce, J. E. In Vivo Protein Interaction Network Identified with a Novel Real-Time Cross-Linked Peptide Identification Strategy. Journal of Proteome Research 2013, 12 (4), 1569–1579. https://doi.org/10.1021/pr3011638.

(17) Tan, D.; Li, Q.; Zhang, M.-J.; Liu, C.; Ma, C.; Zhang, P.; Ding, Y.-H.; Fan, S.-B.; Tao, L.; Yang, B.; et al. Trifunctional Cross-Linker for Mapping Protein-Protein Interaction Networks and Comparing Protein Conformational States. eLife 2016, 5. https://doi.org/10.7554/eLife.12509.

(18) Petrotchenko, E. V.; Serpa, J. J.; Borchers, C. H. An Isotopically Coded CID-Cleavable Biotinylated Cross-Linker for Structural Proteomics. Molecular & Cellular Proteomics 2011, 10 (2), M110.001420–M110.001420. https://doi.org/10.1074/mcp.M110.001420.

(19) Luo, J.; Fishburn, J.; Hahn, S.; Ranish, J. An Integrated Chemical Cross-Linking and Mass Spectrometry Approach to Study Protein Complex Architecture and Function. Mol. Cell Proteomics 2012, 11 (2), M111.008318. https://doi.org/10.1074/mcp.M111.008318.

(20) Petrotchenko, E. V.; Borchers, C. H. Application of a Fast Sorting Algorithm to the Assignment of Mass Spectrometric Cross-Linking Data. Proteomics 2014, 14 (17–18), 1987–1989. https://doi.org/10.1002/pmic.201300486.

(21) Yang, B.; Wu, Y.-J.; Zhu, M.; Fan, S.-B.; Lin, J.; Zhang, K.; Li, S.; Chi, H.; Li, Y.-X.; Chen, H.-F.; et al. Identification of Cross-Linked Peptides from Complex Samples. Nat. Methods 2012, 9 (9), 904–906. https://doi.org/10.1038/nmeth.2099.

(22) Kao, A.; Chiu, C. -l.; Vellucci, D.; Yang, Y.; Patel, V. R.; Guan, S.; Randall, A.; Baldi, P.; Rychnovsky, S. D.; Huang, L. Development of a Novel Cross-Linking Strategy for Fast and Accurate Identification of Cross-Linked Peptides of Protein Complexes. Molecular & Cellular Proteomics 2011, 10 (1), M110.002212–M110.002212. https://doi.org/10.1074/mcp.M110.002212.

(23) Jewett, J. C.; Bertozzi, C. R. Cu-Free Click Cycloaddition Reactions in Chemical Biology. Chem Soc Rev 2010, 39 (4), 1272–1279.

(24) Burke, A. M.; Kandur, W.; Novitsky, E. J.; Kaake, R. M.; Yu, C.; Kao, A.; Vellucci, D.; Huang, L.; Rychnovsky, S. D. Synthesis of Two New Enrichable and MS-Cleavable Cross-Linkers to Define Protein–Protein Interactions by Mass Spectrometry. Org. Biomol. Chem. 2015, 13 (17), 5030–5037. https://doi.org/10.1039/C5OB00488H.

(25) Deng, W.; Shi, X.; Tjian, R.; Lionnet, T.; Singer, R. H. CASFISH: CRISPR/Cas9-Mediated in Situ Labeling of Genomic Loci in Fixed Cells. Proc. Natl. Acad. Sci. U.S.A. 2015, 112 (38), 11870–11875. https://doi.org/10.1073/pnas.1515692112.

(26) Dorfer, V.; Pichler, P.; Stranzl, T.; Stadlmann, J.; Taus, T.; Winkler, S.; Mechtler, K. MS Amanda, a Universal Identification Algorithm Optimized for High Accuracy Tandem Mass Spectra. Journal of Proteome Research 2014, 13 (8), 3679–3684. https://doi.org/10.1021/pr500202e.

(27) Liu, F.; Lössl, P.; Scheltema, R.; Viner, R.; Heck, A. J. R. Optimized Fragmentation Schemes and Data Analysis Strategies for Proteome-Wide Cross-Link Identification. Nature Communications 2017, 8, 15473. https://doi.org/10.1038/ncomms15473.

(28) Chambers, M. C.; Maclean, B.; Burke, R.; Amodei, D.; Ruderman, D. L.; Neumann, S.; Gatto, L.; Fischer, B.; Pratt, B.; Egertson, J.; et al. A Cross-Platform Toolkit for Mass Spectrometry and Proteomics. Nat Biotechnol 2012, 30 (10), 918–920. https://doi.org/10.1038/nbt.2377.

(29) Doblmann, J.; Dusberger, F.; Imre, R.; Hudecz, O.; Stanek, F.; Mechtler, K.; Dürnberger, G. ApQuant: Accurate Label-Free Quantification by Quality Filtering. J. Proteome Res. 2018. https://doi.org/10.1021/acs.jproteome.8b00113.

(30) Perez-Riverol, Y.; Csordas, A.; Bai, J.; Bernal-Llinares, M.; Hewapathirana, S.; Kundu, D. J.; Inuganti, A.; Griss, J.; Mayer, G.; Eisenacher, M.; et al. The PRIDE Database and Related Tools and Resources in 2019: Improving Support for Quantification Data. Nucleic Acids Res. 2019, 47 (D1), D442–D450. https://doi.org/10.1093/nar/gky1106.

(31) Iacobucci, C.; Götze, M.; Ihling, C. H.; Piotrowski, C.; Arlt, C.; Schäfer, M.; Hage, C.; Schmidt, R.; Sinz, A. A Cross-Linking/Mass Spectrometry Workflow Based on MS-Cleavable Cross-Linkers and the MeroX Software for Studying Protein Structures and Protein–Protein Interactions. Nature Protocols 2018, 13 (12), 2864. https://doi.org/10.1038/s41596-018-0068-8.

(32) Stadlmeier, M.; Runtsch, L. S.; Streshnev, F.; Wühr, M.; Carell, T. A Click-Chemistry-Based Enrichable Crosslinker for Structural and Protein Interaction Analysis by Mass Spectrometry. Chembiochem 2020, 21 (1–2), 103–107. https://doi.org/10.1002/cbic.201900611.

(33) Yang, J.; Gupta, V.; Tallman, K. A.; Porter, N. A.; Carroll, K. S.; Liebler, D. C. Global, in Situ, Site-Specific Analysis of Protein S-Sulfenylation. Nat Protoc 2015, 10 (7), 1022–1037. https://doi.org/10.1038/nprot.2015.062.

(34) Nainar, S.; Beasley, S.; Fazio, M.; Kubota, M.; Dai, N.; Corrêa, I. R.; Spitale, R. C. Metabolic Incorporation of Azide Functionality into Cellular RNA. Chembiochem 2016, 17 (22), 2149–2152. https://doi.org/10.1002/cbic.201600300.

