## Supporting Information for "Fast and highly efficient affinity enrichment of Azide-A-DSBSO cross-linked peptides"

### Table of Contents

**Supplemental Figure 1: Estimated enrichment factor.**

**Supplemental Figure 2: Multiple incubation of DSBSO linked Cas9 to DBCO beads.**

**Supplemental Figure 3: Enrichment of *E. coli* ribosomal cross-linked peptides.**

**Supplemental Figure 4: A closer look to the enrichment using cross-linked *E. coli* ribosome.**

**Supplemental Figure 5: Higher protein input is possible after enrichment, leading to more cross-links.**

**Supplemental Figure 6: Error estimation – spiked *E. coli* ribosome samples.**

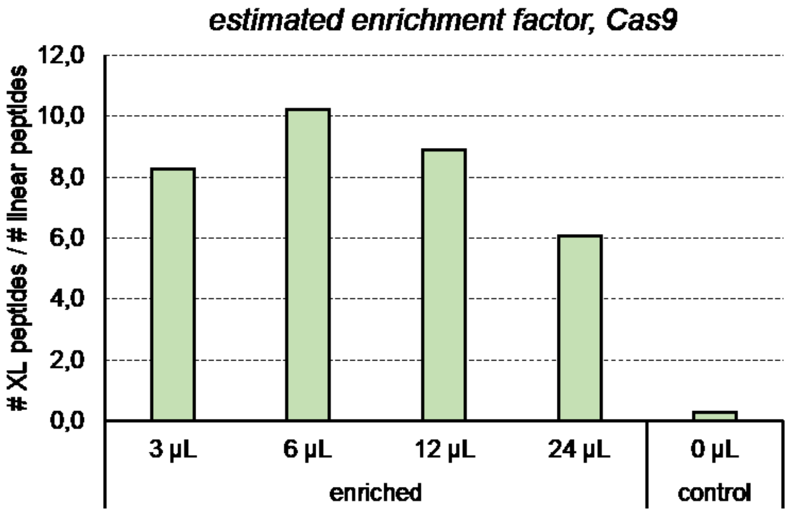


Supplemental Figure 1: **Estimated enrichment factor.** 20 µg Cas9 were cross-linked with 0.5 mM DSBSO. Bars indicate the ratio cross-linked peptides (CSMs via XlinkX)) over linear peptides (PSMs excluding DSBSO modified peptides, via MS Amanda) with or without enrichment (control) using the indicated bead slurry volumes.

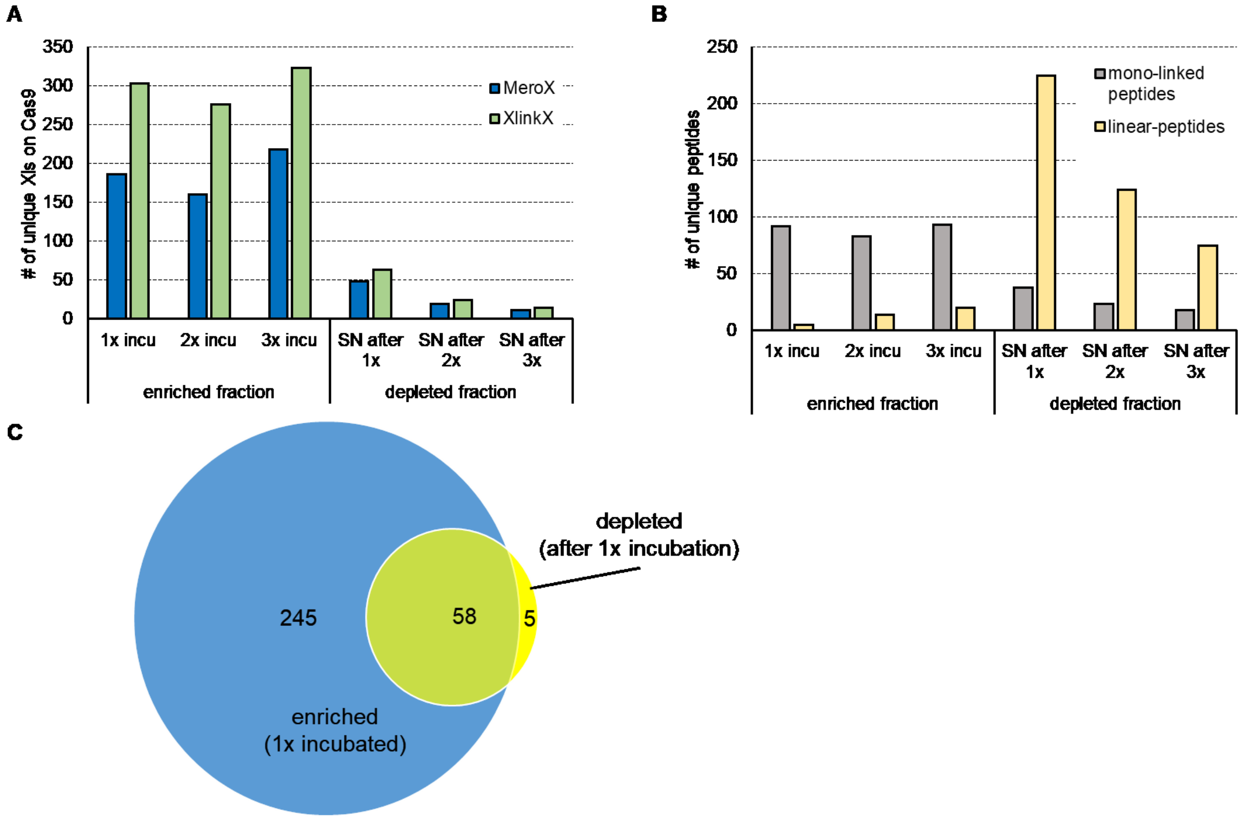


Supplemental Figure 2: **Multiple incubation of DSBSO linked Cas9 to DBCO beads.** 20 µg recombinant Cas9 protein were cross-linked using 0.5 mM DSBSO. After digestion, cross-linked peptides were enriched by incubation to 12 µL DBCO bead slurry, washed and eluted as described under materials and methods (= "1x incu"). Alternatively, the remaining supernatant was again incubated to 12 µL bead slurry (= "2x incu") which was repeated a second time in the last experiment (= "3x incu"). For 2x and 3x incubated sample the resulting DSBSO loaded beads of each step were pooled, washed and cross-linked peptides were eluted as done for the 1x incubated sample respectively. The last remaining supernatant (after 1, 2 or 3x incubation to the beads) was kept and analyzed as depleted fraction. Bars indicate the number of unique cross-links within Cas9 (**A**) or unique mono-linked peptides (as given by MeroX) and linear peptides (as given by MS Amanda) (**B**). The Venn diagram shows the overlap of detected unique cross-links obtained after a single incubation to DBCO beads (enriched fraction) compared to the remaining supernatant over the beads (depleted fraction) (**C**). Results filtered for 5 % FDR (Cross-links, mono links) or 1 % FDR (linear peptides), n=1

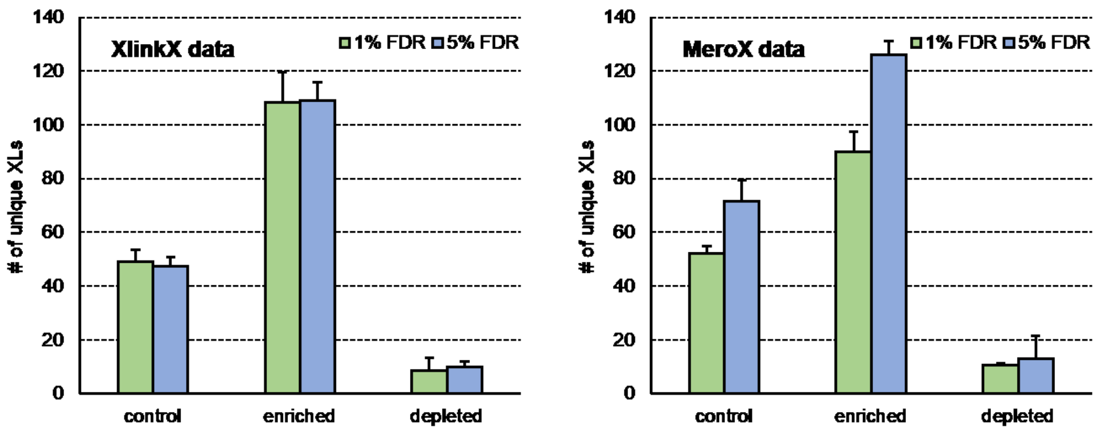


Supplemental Figure 3**: Enrichment of *E. coli* ribosomal cross-linked peptides.** Number of unique cross-links detected within ribosome-shotgun database after DBCO bead enrichment (enriched) or in the remaining supernatant over the beads after click reaction (depleted). 20 µg *E. coli* ribosome each were linked using 1 mM DSBSO. Bars indicate the average values, with standard deviation depicted as error bar. No additional score cutoff used for MeroX, standard score cutoff of 40 for XlinkX used, FDR and search engine as indicated, n≥3.

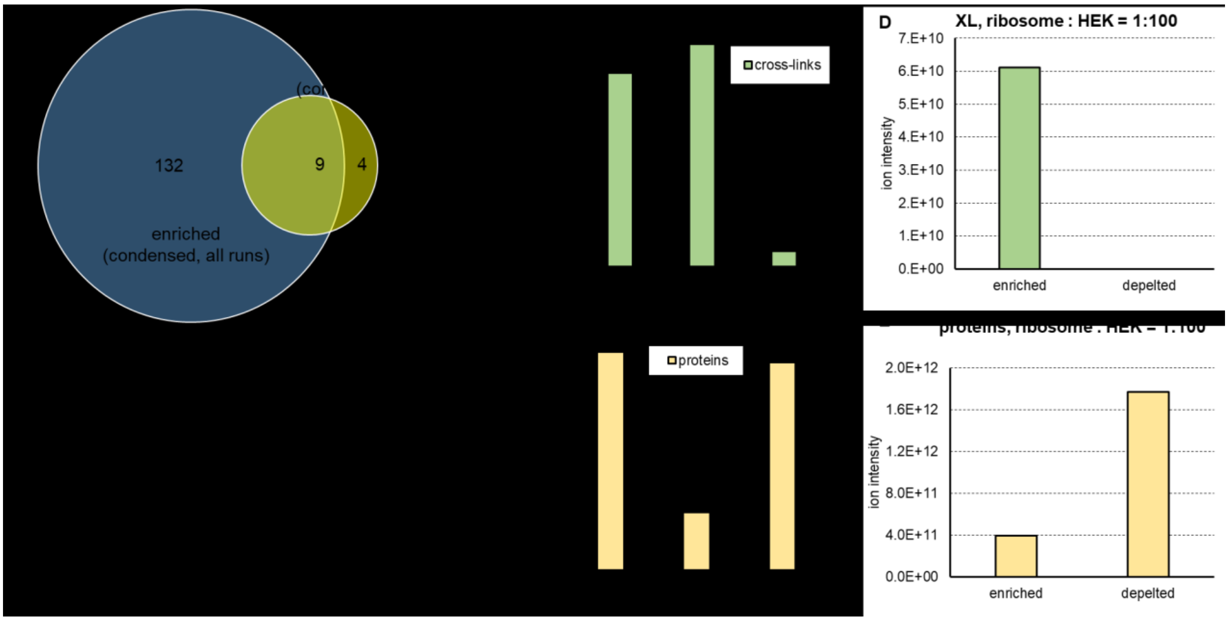


Supplemental Figure 4: **A closer look to the enrichment using cross-linked *E. coli* ribosome.** This Figure contains the experimental data shown in Figure 3 in panel A-C as well as data from Figure 4 in panel D-E. (**A**) The Venn diagram shows the overlap of all unique-cross links of enriched vs depleted fractions found in 3 replicates. (**B, C**) The bar plots show the ion intensities ± SD of cross-links (**B**) or proteins (**C**) after LFQ in the indicated fractions in unspiked ribosome, average values, cross-links as detected using XlinkX, proteins as detected with MS Amanda, n=3. (**D, E**)**:** The bar plots show the ion intensities ± SD of cross-links (**D**) or proteins (**E**) after LFQ in the indicated fractions in ribosome spiked in a HEK background (1:100), cross-links as detected using XlinkX, proteins as detected with MS Amanda.

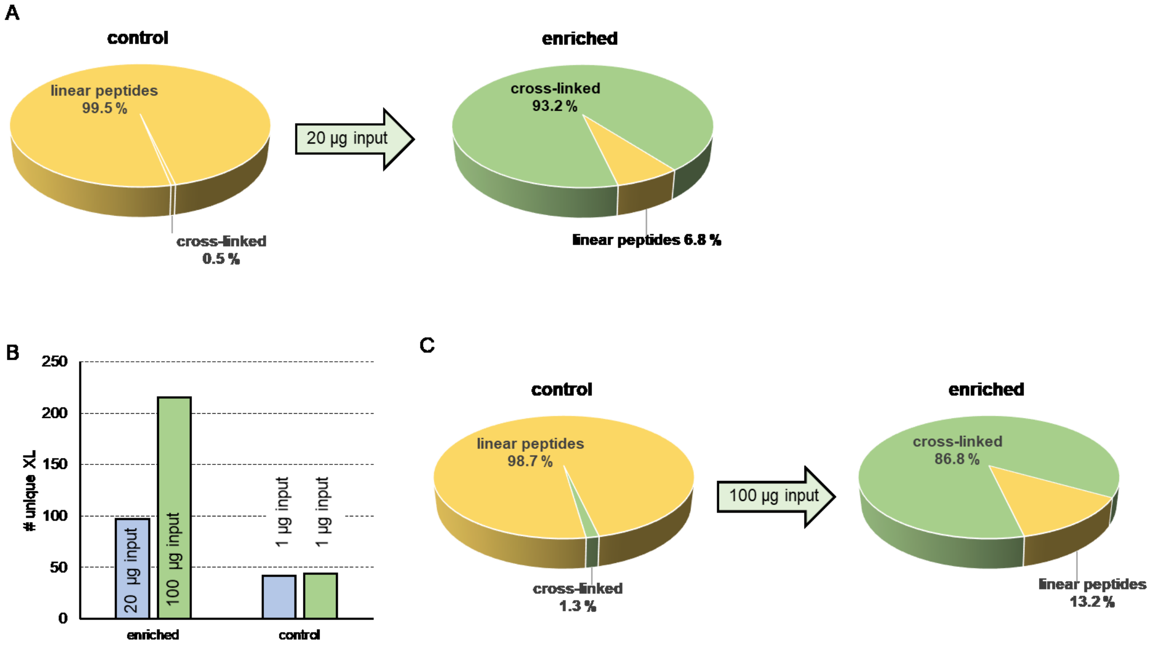


Supplemental Figure 5: **Higher protein input is possible after enrichment, leading to more cross-links.** 20 µg *E. coli* ribosome were cross-linked and enriched using DBCO beads. Injection of 1 µg total protein (without enrichment, control) or 100 % of the eluate (enriched), relative ion intensities shown after LFQ, average values, n=3 (**A**), The same experiment was repeated using 100 µg of input material for enrichment, unique cross-link numbers (XlinkX, 5% FDR) are shown in (**B**), relative ion intensities after LFQ in (**C**).

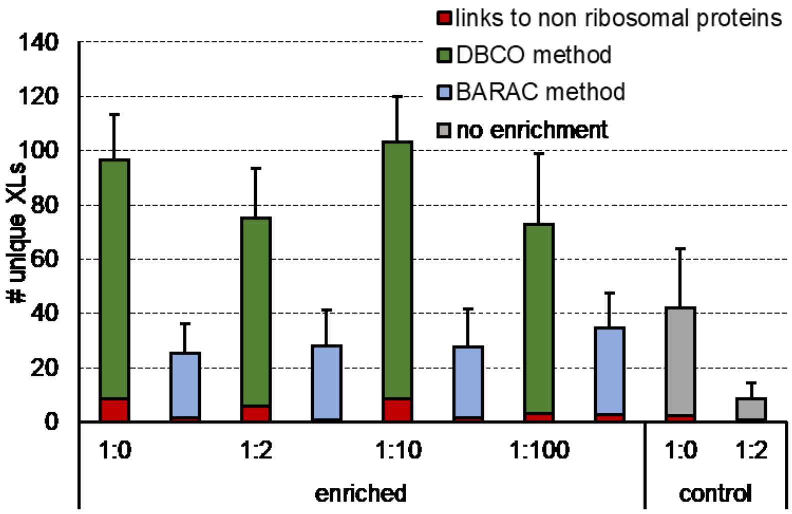


Supplemental Figure 6: **Error estimation – spiked *E. coli* ribosome samples.** Found average number of unique cross-links from the recovery experiment, when analyzing data via XlinkX using a combined human Swiss-Prot + a shotgun database originating from *E. coli* ribosome. Hits within this shotgun database are potential true cross-links (indicated in green or blue respectively), while hits within the human proteome should not exist (indicated in red).

Supplementary Table 1: List of detected unique DSBSO cross-links given by XlinkX in enriched and depleted fraction of cross-linked *E. coli* ribosome (Data from Figure 3 / Supplemental Figure 4)

| in enriched | in depleted | Max. XlinkX Score | Crosslink Type | # CSMs | Sequence A | Modifications A | Accession A | Position A | Sequence B | Modifications B | Accession B | Position B |
|---|---|---|---|---|---|---|---|---|---|---|---|---|
| y | y | 187.45 | Intra | 16 | AIAEQL[K]YTGNK | 1xDSBSO_linker [K7] | P0C018 | 63 | DAAAAVG[K]AVAER | 1xDSBSO_linker [K8] | P0C018 | 76 |
| y | y | 169.69 | Intra | 18 | [K]ALEEAGAEVEVK | 1xDSBSO_linker [K1] | P0A7K2 | 109 | EGVS[K]DDAEALK | 1xDSBSO_linker [K5] | P0A7K2 | 101 |
| y | y | 142.27 | Intra | 6 | TEFDVIL[K]AAGANK | 1xDSBSO_linker [K8] | P0A7K2 | 60 | DDAEAL[K]K | 1xDSBSO_linker [K7] | P0A7K2 | 108 |
| y | y | 130.84 | Intra | 10 | DAAAAVG[K]AVAER | 1xDSBSO_linker [K8] | P0C018 | 76 | GI[K]DVSFDR | 1xDSBSO_linker [K3] | P0C018 | 88 |
| y | y | 126.6 | Intra | 61 | EA[K]DLVESAPAALK | 1xDSBSO_linker [K3] | P0A7K2 | 85 | VAVI[K]AVR | 1xDSBSO_linker [K5] | P0A7K2 | 71 |
| y | y | 122.17 | Intra | 7 | VGTVTPNVAEAV[K]NAK | 1xDSBSO_linker [K13] | P0A7L0 | 154 | VDFDAD[K]LK | 1xDSBSO_linker [K7] | P0A7L0 | 184 |
| y | y | 112.7 | Intra | 78 | [K]PELDAK | 1xDSBSO_linker [K1] | P0A7V3 | 108 | LGA[K]GIK | 1xDSBSO_linker [K4] | P0A7V3 | 147 |
| y | y | 91.44 | Inter | 9 | RSHDALTAVTSLSVD[K]TSGEK | 1xDSBSO_linker [K16] | P0A7N4 | 32 | SE[K]AEAAAE | 1xDSBSO_linker [K3] | P0AG44 | 121 |
| y | y | 43.67 | Intra | 2 | VGFFNPIASE[K]EEGTR | 1xDSBSO_linker [K11] | P0A7T3 | 46 | EVN[K]AA | 1xDSBSO_linker [K4] | P0A7T3 | 80 |
| n | y | 132.9 | Intra | 3 | LVDIVEPTE[K]TVDALMR | 1xOxidation [M16]; 1xDSBSO_linker [K10] | P0A7R5 | 82 | LIDQATAEIVETA[K]R | 1xDSBSO_linker [K14] | P0A7R5 | 30 |
| n | y | 114.64 | Intra | 1 | SG[K]SELEAFEVALENVRPTVEVK | 1xDSBSO_linker [K3] | P02359 | 56 | LANELSDAAEN[K]GTAVK | 1xDSBSO_linker [K12] | P02359 | 131 |
| n | y | 122.76 | Inter | 40 | AGDEG[K]LFGSIGTR | 1xDSBSO_linker [K6] | P0A7R1 | 89 | [K]VEADCR | 1xCarbamidomethyl [C6]; 1xDSBSO_linker [K1] | P60422 | 183 |
| n | y | 109.6 | Intra | 1 | ALEEAGAEVEV[K] | 1xDSBSO_linker [K12] | P0A7K2 | 121 | ALEEAGAEVEV[K] | 1xDSBSO_linker [K7] | P0A7K2 | 108 |
| y | n | 167.98 | Intra | 12 | AAAQ[K]AFNEMQPIVDR | 1xDSBSO_linker [K5] | P0A7U7 | 49 | [K]VYAAIEAGDK | 1xDSBSO_linker [K1] | P0A7U7 | 34 |
| y | n | 161.49 | Inter | 19 | [K]LQLVGVGYR | 1xDSBSO_linker [K1] | P0AG55 | 86 | VAVI[K]AVR | 1xDSBSO_linker [K5] | P0A7K2 | 71 |
| y | n | 150.27 | Intra | 5 | GLMPNP[K]VGTVTPNVAEAVK | 1xDSBSO_linker [K7] | P0A7L0 | 141 | ND[K]NGIIHTTIGK | 1xDSBSO_linker [K3] | P0A7L0 | 167 |
| y | n | 145.21 | Intra | 26 | [K]ALEEAGAEVEVK | 1xDSBSO_linker [K1] | P0A7K2 | 109 | VAVI[K]AVR | 1xDSBSO_linker [K5] | P0A7K2 | 71 |
| y | n | 141.48 | Intra | 26 | GATGLGL[K]EAK | 1xDSBSO_linker [K8] | P0A7K2 | 82 | VAVI[K]AVR | 1xDSBSO_linker [K5] | P0A7K2 | 71 |
| y | n | 140.98 | Intra | 24 | AAGI[K]SGSGKPNK | 1xDSBSO_linker [K5] | P0A7J7 | 87 | TPPAAVLL[K]K | 1xDSBSO_linker [K9] | P0A7J7 | 81 |
| y | n | 137.83 | Intra | 33 | GVTVD[K]MTELRK | 1xDSBSO_linker [K6] | P0A7J3 | 37 | ANA[K]FEVK | 1xDSBSO_linker [K4] | P0A7J3 | 105 |
| y | n | 137.5 | Intra | 10 | VFQTHSPVVDSISV[K]R | 1xDSBSO_linker [K15] | P0A7K6 | 87 | [K]ISNGEGVER | 1xDSBSO_linker [K1] | P0A7K6 | 63 |
| y | n | 135.84 | Intra | 9 | EA[K]DLVESAPAALK | 1xDSBSO_linker [K3] | P0A7K2 | 85 | AAGAN[K]VAVIK | 1xDSBSO_linker [K6] | P0A7K2 | 66 |

| | | | | | | | | | | | | |
|---|---|---|---|---|---|---|---|---|---|---|---|---|
| y | n | 128.27 | Intra | 3 | TLNDAVEV[K]HADNTLTFGPR | 1xDSBSO_linker [K9] | P0AG55 | 44 | VA[K]APVVVPAGVDVK | 1xDSBSO_linker [K3] | P0AG55 | 6 |
| y | n | 127.53 | Intra | 22 | AKPTQA[K]GVYIK | 1xDSBSO_linker [K7] | P0A7L0 | 205 | [K]SDQNVR | 1xDSBSO_linker [K1] | P0A7L0 | 54 |
| y | n | 124.34 | Intra | 22 | A[K]DEADEKDAIATVNK | 1xDSBSO_linker [K2] | P0AG67 | 522 | [K]GAIVTGK | 1xDSBSO_linker [K1] | P0AG67 | 450 |
| y | n | 123.53 | Inter | 5 | ANA[K]FEVK | 1xDSBSO_linker [K4] | P0A7J3 | 105 | VAVI[K]AVR | 1xDSBSO_linker [K5] | P0A7K2 | 71 |
| y | n | 120.76 | Intra | 15 | VTIHTARPGIVIG[K]K | 1xDSBSO_linker [K14] | P0A7V3 | 79 | ELA[K]ASVSR | 1xDSBSO_linker [K4] | P0A7V3 | 49 |
| y | n | 120.76 | Intra | 7 | L[K]DLETQSQDGTFDK | 1xDSBSO_linker [K2] | P0A7V0 | 115 | G[K]ILFVGTK | 1xDSBSO_linker [K2] | P0A7V0 | 66 |
| y | n | 119.58 | Intra | 6 | V[K]GGFTVELNGIR | 1xDSBSO_linker [K2] | P0AG67 | 117 | VI[K]LDQK | 1xDSBSO_linker [K3] | P0AG67 | 158 |
| y | n | 118.18 | Intra | 6 | VT[K]IFVDEGPSMK | 1xDSBSO_linker [K3] | P61175 | 73 | METIA[K]HR | 1xDSBSO_linker [K6] | P61175 | 6 |
| y | n | 115.01 | Intra | 4 | AKDEADE[K]DAIATVNK | 1xDSBSO_linker [K8] | P0AG67 | 528 | [K]GAIVTGK | 1xDSBSO_linker [K1] | P0AG67 | 450 |
| y | n | 114.59 | Intra | 3 | GATGLGL[K]EAK | 1xDSBSO_linker [K8] | P0A7K2 | 82 | AAGAN[K]VAVIK | 1xDSBSO_linker [K6] | P0A7K2 | 66 |
| y | n | 113.5 | Inter | 5 | EA[K]DLVESAPAALK | 1xDSBSO_linker [K3] | P0A7K2 | 85 | [K]LQLVGVGYR | 1xDSBSO_linker [K1] | P0AG55 | 86 |
| y | n | 112.7 | Inter | 38 | [K]ILPDPK | 1xDSBSO_linker [K1] | P02359 | 11 | [K]AGFVTR | 1xDSBSO_linker [K1] | P0A7X3 | 100 |
| y | n | 112.31 | Intra | 12 | VFQTHSPVVDSISV[K]R | 1xDSBSO_linker [K15] | P0A7K6 | 87 | TG[K]AAR | 1xDSBSO_linker [K3] | P0A7K6 | 106 |
| y | n | 112.31 | Inter | 18 | DIATL[K]NYITESGK | 1xDSBSO_linker [K6] | P0A7T7 | 30 | PVI[K]VR | 1xDSBSO_linker [K4] | P68679 | 4 |
| y | n | 111.69 | Intra | 7 | KGPFIDLHLL[K]K | 1xDSBSO_linker [K11] | P0A7U3 | 17 | VE[K]AVESGDK | 1xDSBSO_linker [K3] | P0A7U3 | 21 |
| y | n | 110.43 | Inter | 22 | VVSD[K]MEK | 1xDSBSO_linker [K5] | P0AG63 | 16 | LLDYL[K]R | 1xDSBSO_linker [K6] | P0ADZ4 | 71 |
| y | n | 109.53 | Intra | 3 | GEILGGMAAVEQPE[K]PAAQPK | 1xDSBSO_linker [K15] | P0A7V3 | 219 | [K]PELDAK | 1xDSBSO_linker [K1] | P0A7V3 | 108 |
| y | n | 108.98 | Inter | 3 | INGQVITI[K]GK | 1xDSBSO_linker [K9] | P0AG55 | 27 | VAVI[K]AVR | 1xDSBSO_linker [K5] | P0A7K2 | 71 |
| y | n | 107.73 | Intra | 15 | AIQSE[K]AR | 1xDSBSO_linker [K6] | P0A7U7 | 16 | [K]HNASR | 1xDSBSO_linker [K1] | P0A7U7 | 19 |
| y | n | 107.73 | Intra | 4 | DDAEAL[K]K | 1xDSBSO_linker [K7] | P0A7K2 | 108 | VAVI[K]AVR | 1xDSBSO_linker [K5] | P0A7K2 | 71 |
| y | n | 107.73 | Inter | 6 | [K]NIEFFEAR | 1xDSBSO_linker [K1] | P0A7R1 | 42 | FWVESE[K]R | 1xDSBSO_linker [K7] | P0A7M2 | 44 |
| y | n | 107.73 | Intra | 1 | ITLNMGVGEAIAD[K]K | 1xDSBSO_linker [K14] | P62399 | 47 | SVAGF[K]IR | 1xDSBSO_linker [K6] | P62399 | 78 |
| y | n | 107.15 | Inter | 6 | GLSA[K]SFDGR | 1xDSBSO_linker [K5] | P62399 | 120 | EISMSI[K]R | 1xDSBSO_linker [K7] | P0A7S9 | 78 |
| y | n | 107.15 | Inter | 7 | SVAGF[K]IR | 1xDSBSO_linker [K6] | P62399 | 78 | GHAAD[K]K | 1xDSBSO_linker [K6] | P0A7U3 | 87 |
| y | n | 106.65 | Intra | 38 | YRND[K]NGIIHTTIGK | 1xDSBSO_linker [K5] | P0A7L0 | 167 | [K]SDQNVR | 1xDSBSO_linker [K1] | P0A7L0 | 54 |
| y | n | 104.94 | Intra | 9 | I[K]LVSSAGTGHFYTTTK | 1xDSBSO_linker [K2] | P0A7N9 | 10 | TKPE[K]LELK | 1xDSBSO_linker [K5] | P0A7N9 | 33 |
| y | n | 104.64 | Intra | 5 | VI[K]LDQK | 1xDSBSO_linker [K3] | P0AG67 | 158 | VL[K]FDR | 1xDSBSO_linker [K3] | P0AG67 | 247 |

| | | | | | | | | | | | | |
|---|---|---|---|---|---|---|---|---|---|---|---|---|
| y | n | 104.15 | Inter | 22 | G[K]NGELTR | 1xDSBSO_linker [K2] | P0AG55 | 29 | VAVI[K]AVR | 1xDSBSO_linker [K5] | P0A7K2 | 71 |
| y | n | 101.72 | Inter | 8 | [K]VSQALDILTYTNK | 1xDSBSO_linker [K1] | P61175 | 28 | TSGE[K]HLR | 1xDSBSO_linker [K5] | P0A7N4 | 37 |
| y | n | 101.72 | Inter | 3 | [K]LQLVGVGYR | 1xDSBSO_linker [K1] | P0AG55 | 86 | GATGLGL[K]EAK | 1xDSBSO_linker [K8] | P0A7K2 | 82 |
| y | n | 98.51 | Intra | 23 | QHVIY[K]EAK | 1xDSBSO_linker [K6] | P0A7N9 | 50 | TKPE[K]LELK | 1xDSBSO_linker [K5] | P0A7N9 | 33 |
| y | n | 96.12 | Intra | 7 | VDFDAD[K]LK | 1xDSBSO_linker [K7] | P0A7L0 | 184 | NA[K]AGQVR | 1xDSBSO_linker [K3] | P0A7L0 | 157 |
| y | n | 95.35 | Intra | 2 | A[K]LHDYYK | 1xDSBSO_linker [K2] | P62399 | 2 | DEVV[K]K | 1xDSBSO_linker [K5] | P62399 | 14 |
| y | n | 95.35 | Intra | 2 | DTLHLEG[K]ELEFK | 1xDSBSO_linker [K8] | P0AG67 | 150 | VL[K]FDR | 1xDSBSO_linker [K3] | P0AG67 | 247 |
| y | n | 94.27 | Intra | 14 | SGSGKPN[K]DK | 1xDSBSO_linker [K8] | P0A7J7 | 95 | VG[K]ISR | 1xDSBSO_linker [K3] | P0A7J7 | 100 |
| y | n | 92.15 | Inter | 10 | VAVI[K]AVR | 1xDSBSO_linker [K5] | P0A7K2 | 71 | [K]AAGIK | 1xDSBSO_linker [K1] | P0A7J7 | 82 |
| y | n | 92.08 | Intra | 3 | AAGAN[K]VAVIK | 1xDSBSO_linker [K6] | P0A7K2 | 66 | VAVI[K]AVR | 1xDSBSO_linker [K5] | P0A7K2 | 71 |
| y | n | 91.93 | Intra | 2 | LIVVE[K]FSVEAPK | 1xDSBSO_linker [K6] | P60723 | 123 | LLAQ[K]LK | 1xDSBSO_linker [K5] | P60723 | 137 |
| y | n | 91.09 | Intra | 6 | I[K]LVSSAGTGHFYTTK | 1xDSBSO_linker [K2] | P0A7N9 | 10 | QHVIY[K]EAK | 1xDSBSO_linker [K6] | P0A7N9 | 50 |
| y | n | 89.75 | Intra | 5 | QNGISYS[K]FINGLK | 1xDSBSO_linker [K8] | P0A7L3 | 78 | [K]ASVEIDRK | 1xDSBSO_linker [K1] | P0A7L3 | 85 |
| y | n | 89.08 | Inter | 3 | LIDQATAEIVETA[K]R | 1xDSBSO_linker [K14] | P0A7R5 | 30 | [K]GAIVTGK | 1xDSBSO_linker [K1] | P0AG67 | 450 |
| y | n | 87.26 | Intra | 20 | [K]ISNGEGVER | 1xDSBSO_linker [K1] | P0A7K6 | 63 | TG[K]AAR | 1xDSBSO_linker [K3] | P0A7K6 | 106 |
| y | n | 87.26 | Inter | 9 | LPI[K]TTFVTK | 1xDSBSO_linker [K4] | P0ADY7 | 127 | HPY[K]PK | 1xDSBSO_linker [K4] | P68919 | 83 |
| y | n | 87.26 | Inter | 2 | VL[K]FDR | 1xDSBSO_linker [K3] | P0AG67 | 247 | PVI[K]VR | 1xDSBSO_linker [K4] | P68679 | 4 |
| y | n | 86.63 | Inter | 2 | LF[K]EFAK | 1xDSBSO_linker [K3] | P0A7J3 | 97 | VAVI[K]AVR | 1xDSBSO_linker [K5] | P0A7K2 | 71 |
| y | n | 86.34 | Intra | 8 | VAFTALVE[K]AK | 1xDSBSO_linker [K9] | P0A7L3 | 112 | [K]ASVEIDRK | 1xDSBSO_linker [K1] | P0A7L3 | 85 |
| y | n | 85.33 | Inter | 2 | VA[K]APVVVPAGVDVK | 1xDSBSO_linker [K3] | P0AG55 | 6 | VAVI[K]AVR | 1xDSBSO_linker [K5] | P0A7K2 | 71 |
| y | n | 85.06 | Inter | 3 | GLMPNP[K]VGTVTPNVAEAVK | 1xDSBSO_linker [K7] | P0A7L0 | 141 | QHVIY[K]EAK | 1xDSBSO_linker [K6] | P0A7N9 | 50 |
| y | n | 84.61 | Intra | 14 | GLMPNP[K]VGTVTPNVAEAVK | 1xDSBSO_linker [K7] | P0A7L0 | 141 | [K]AKPTQAK | 1xDSBSO_linker [K1] | P0A7L0 | 198 |
| y | n | 83.7 | Intra | 2 | EGVS[K]DDAEALK | 1xDSBSO_linker [K5] | P0A7K2 | 101 | VAVI[K]AVR | 1xDSBSO_linker [K5] | P0A7K2 | 71 |
| y | n | 82.63 | Inter | 7 | VAVI[K]AVR | 1xDSBSO_linker [K5] | P0A7K2 | 71 | VG[K]ISR | 1xDSBSO_linker [K3] | P0A7J7 | 100 |
| y | n | 81.22 | Inter | 6 | [K]GAIVTGK | 1xDSBSO_linker [K1] | P0AG67 | 450 | LGA[K]GIK | 1xDSBSO_linker [K4] | P0A7V3 | 147 |
| y | n | 81 | Intra | 7 | E[K]PTWLEVDAGK | 1xDSBSO_linker [K2] | P0A7V8 | 167 | MEGTF[K]R | 1xDSBSO_linker [K6] | P0A7V8 | 183 |
| y | n | 79.45 | Intra | 2 | VT[K]PEAGHFAK | 1xDSBSO_linker [K3] | P60438 | 62 | AIQVTTGA[K]K | 1xDSBSO_linker [K9] | P60438 | 55 |
| y | n | 79.12 | Inter | 2 | EGVS[K]DDAEALK | 1xDSBSO_linker [K5] | P0A7K2 | 101 | LF[K]EFAK | 1xDSBSO_linker [K3] | P0A7J3 | 97 |
| y | n | 78.19 | Intra | 10 | [K]SDQNVR | 1xDSBSO_linker [K1] | P0A7L0 | 54 | GVYI[K]K | 1xDSBSO_linker [K5] | P0A7L0 | 210 |

| | | | | | | | | | | | |
|---|---|---|---|---|---|---|---|---|---|---|---|
| y | n | 78.19 | Intra | 6 | [K]WLEER | 1xDSBSO_linker [K1] | P38521 | 87 | [K]LDEVR | 1xDSBSO_linker [K1] | P38521 | 93 |
| y | n | 78.16 | Intra | 1 | GEILGGMAAVEQPE[K]PAAQPK | 1xDSBSO_linker [K15] | P0A7V3 | 219 | LGA[K]GIK | 1xDSBSO_linker [K4] | P0A7V3 | 147 |
| y | n | 77.93 | Intra | 2 | ND[K]NGIIHTTIGK | 1xDSBSO_linker [K3] | P0A7L0 | 167 | GVYI[K]K | 1xDSBSO_linker [K5] | P0A7L0 | 210 |
| y | n | 77.72 | Inter | 4 | GLMPNP[K]VGTVTPNVAEAVK | 1xDSBSO_linker [K7] | P0A7L0 | 141 | TKPE[K]LELK | 1xDSBSO_linker [K5] | P0A7N9 | 33 |
| y | n | 77.53 | Inter | 4 | AKPTQA[K]GVYIK | 1xDSBSO_linker [K7] | P0A7L0 | 205 | EA[K]IK | 1xDSBSO_linker [K3] | P0A7N9 | 53 |
| y | n | 77.2 | Intra | 4 | FSVEAP[K]TK | 1xDSBSO_linker [K7] | P60723 | 130 | LLAQ[K]LK | 1xDSBSO_linker [K5] | P60723 | 137 |
| y | n | 76.42 | Intra | 10 | AAN[K]FPAIIYGGK | 1xDSBSO_linker [K4] | P68919 | 25 | [K]EQGK | 1xDSBSO_linker [K1] | P68919 | 10 |
| y | n | 76.27 | Intra | 6 | SGSG[K]PNKDK | 1xDSBSO_linker [K5] | P0A7J7 | 92 | VG[K]ISR | 1xDSBSO_linker [K3] | P0A7J7 | 100 |
| y | n | 73.37 | Inter | 1 | GHAAD[K]K | 1xDSBSO_linker [K6] | P0A7U3 | 87 | [K]PIKK | 1xDSBSO_linker [K1] | P0A7S9 | 114 |
| y | n | 72.62 | Intra | 1 | ANPWQQFAETHN[K]GDRVEGK | 1xDSBSO_linker [K13] | P0AG67 | 363 | [K]GAIVTGK | 1xDSBSO_linker [K1] | P0AG67 | 450 |
| y | n | 72.34 | Inter | 5 | DLETQSQDGTFD[K]LTK | 1xDSBSO_linker [K13] | P0A7V0 | 128 | LGA[K]GIK | 1xDSBSO_linker [K4] | P0A7V3 | 147 |
| y | n | 72.34 | Inter | 6 | GLSA[K]SFDGR | 1xDSBSO_linker [K5] | P62399 | 120 | RFNIPGS[K] | 1xDSBSO_linker [K8] | P0A7M9 | 70 |
| y | n | 72 | Intra | 3 | GYLDYDA[K]K | 1xDSBSO_linker [K8] | P07012 | 30 | AQALG[K]ER | 1xDSBSO_linker [K6] | P07012 | 58 |
| y | n | 72 | Inter | 2 | GVTVD[K]MTELR | 1xDSBSO_linker [K6] | P0A7J3 | 37 | VAVI[K]AVR | 1xDSBSO_linker [K5] | P0A7K2 | 71 |
| y | n | 71.07 | Intra | 5 | VGTVTPNVAEAV[K]NAK | 1xDSBSO_linker [K13] | P0A7L0 | 154 | [K]AKPTQAK | 1xDSBSO_linker [K1] | P0A7L0 | 198 |
| y | n | 71.07 | Inter | 4 | QHVIY[K]EAK | 1xDSBSO_linker [K6] | P0A7N9 | 50 | [K]AKPTQAK | 1xDSBSO_linker [K1] | P0A7L0 | 198 |
| y | n | 71.07 | Intra | 3 | VTIHTARPGIVIG[K]K | 1xDSBSO_linker [K14] | P0A7V3 | 79 | GEDVE[K]LR | 1xDSBSO_linker [K6] | P0A7V3 | 86 |
| y | n | 70.86 | Inter | 2 | GHAAD[K]K | 1xDSBSO_linker [K6] | P0A7U3 | 87 | KPIK[K] | 1xDSBSO_linker [K5] | P0A7S9 | 118 |
| y | n | 69.2 | Intra | 2 | AAGI[K]SGSGKPNK | 1xDSBSO_linker [K5] | P0A7J7 | 87 | VG[K]ISR | 1xDSBSO_linker [K3] | P0A7J7 | 100 |
| y | n | 68.6 | Intra | 14 | RTKPE[K]LELK | 1xDSBSO_linker [K6] | P0A7N9 | 33 | EA[K]IK | 1xDSBSO_linker [K3] | P0A7N9 | 53 |
| y | n | 68.6 | Intra | 10 | TLLNE[K]AGA | 1xDSBSO_linker [K6] | P0A7M6 | 60 | A[K]ELR | 1xDSBSO_linker [K2] | P0A7M6 | 4 |
| y | n | 67.76 | Inter | 4 | V[K]TLLNEK | 1xDSBSO_linker [K2] | P0A7M6 | 54 | LL[K]VLR | 1xDSBSO_linker [K3] | P0ADZ0 | 9 |
| y | n | 65.44 | Inter | 6 | IVIERPA[K]SIR | 1xDSBSO_linker [K8] | P0A7V3 | 62 | AA[K]GE | 1xDSBSO_linker [K3] | P0AG67 | 555 |
| y | n | 65.33 | Intra | 5 | GVTVD[K]MTELRK | 1xDSBSO_linker [K6] | P0A7J3 | 37 | EFA[K]ANAK | 1xDSBSO_linker [K4] | P0A7J3 | 101 |
| y | n | 65.33 | Intra | 1 | ANA[K]FEVK | 1xDSBSO_linker [K4] | P0A7J3 | 105 | LF[K]EFAK | 1xDSBSO_linker [K3] | P0A7J3 | 97 |
| y | n | 65.26 | Intra | 2 | LANELSDAAEN[K]GTAVK | 1xDSBSO_linker [K12] | P02359 | 131 | RGD[K]SMALR | 1xDSBSO_linker [K4] | P02359 | 114 |
| y | n | 65.26 | Intra | 2 | LPI[K]TTFVTK | 1xDSBSO_linker [K4] | P0ADY7 | 127 | EAF[K]LAAK | 1xDSBSO_linker [K4] | P0ADY7 | 118 |
| y | n | 64.4 | Inter | 2 | A[K]LHDYYK | 1xDSBSO_linker [K2] | P62399 | 2 | FNIPGS[K] | 1xDSBSO_linker [K7] | P0A7M9 | 70 |

| | | | | | | | | | | | |
|---|---|---|---|---|---|---|---|---|---|---|---|
| y | n | 64.4 | Intra | 2 | GLMPNP[K]VGTVTPNVAEAVK | 1xDSBSO_linker [K7] | P0A7L0 | 141 | [K]SDQNVR | 1xDSBSO_linker [K1] | P0A7L0 | 54 |
| y | n | 64.21 | Inter | 1 | EA[K]DLVESAPAALK | 1xDSBSO_linker [K3] | P0A7K2 | 85 | [K]AAGIK | 1xDSBSO_linker [K1] | P0A7J7 | 82 |
| y | n | 63.05 | Inter | 4 | PVI[K]VR | 1xDSBSO_linker [K4] | P68679 | 4 | A[K]DER | 1xDSBSO_linker [K2] | P02358 | 106 |
| y | n | 62.09 | Intra | 3 | SNSETI[K] | 1xDSBSO_linker [K7] | P60624 | 104 | FEDG[K]K | 1xDSBSO_linker [K5] | P60624 | 91 |
| y | n | 61.3 | Inter | 17 | LGA[K]GIK | 1xDSBSO_linker [K4] | P0A7V3 | 147 | QSI[K]R | 1xDSBSO_linker [K4] | P0A7V0 | 112 |
| y | n | 61.12 | Intra | 1 | LETIEGS[K]GK | 1xDSBSO_linker [K8] | P0AFG8 | 724 | [K]VVADAIAK | 1xDSBSO_linker [K1] | P0AFG8 | 866 |
| y | n | 60.87 | Inter | 20 | GHAAD[K]K | 1xDSBSO_linker [K6] | P0A7U3 | 87 | KPI[K]K | 1xDSBSO_linker [K4] | P0A7S9 | 117 |
| y | n | 58.55 | Inter | 1 | SVAGF[K]IR | 1xDSBSO_linker [K6] | P62399 | 78 | FNIPGS[K] | 1xDSBSO_linker [K7] | P0A7M9 | 70 |
| y | n | 58.11 | Intra | 4 | QHVIY[K]EAK | 1xDSBSO_linker [K6] | P0A7N9 | 50 | LEL[K]K | 1xDSBSO_linker [K4] | P0A7N9 | 37 |
| y | n | 55.6 | Intra | 3 | I[K]LVSSAGTGHFYTTTK | 1xDSBSO_linker [K2] | P0A7N9 | 10 | EA[K]IK | 1xDSBSO_linker [K3] | P0A7N9 | 53 |
| y | n | 55.42 | Inter | 4 | DIATL[K]NYITESGK | 1xDSBSO_linker [K6] | P0A7T7 | 30 | A[K]DER | 1xDSBSO_linker [K2] | P02358 | 106 |
| y | n | 54.93 | Inter | 1 | LLDYL[K]R | 1xDSBSO_linker [K6] | P0ADZ4 | 71 | VVE[K]AVL | 1xDSBSO_linker [K4] | P0AG63 | 81 |
| y | n | 53.48 | Intra | 9 | RT[K]PEKLELK | 1xDSBSO_linker [K3] | P0A7N9 | 30 | EA[K]IK | 1xDSBSO_linker [K3] | P0A7N9 | 53 |
| y | n | 53.48 | Intra | 4 | L[K]DLETQSQDGTFDK | 1xDSBSO_linker [K2] | P0A7V0 | 115 | QSI[K]R | 1xDSBSO_linker [K4] | P0A7V0 | 112 |
| y | n | 53.48 | Inter | 3 | VI[K]LDQK | 1xDSBSO_linker [K3] | P0AG67 | 158 | A[K]DER | 1xDSBSO_linker [K2] | P02358 | 106 |
| y | n | 52.59 | Inter | 2 | TSGE[K]HLR | 1xDSBSO_linker [K5] | P0A7N4 | 37 | AAVLV[K]K | 1xDSBSO_linker [K6] | P61175 | 48 |
| y | n | 52.58 | Inter | 1 | V[K]HPSEIVNVGDEITVK | 1xDSBSO_linker [K2] | P0AG67 | 229 | PVI[K]VR | 1xDSBSO_linker [K4] | P68679 | 4 |
| y | n | 52.33 | Intra | 2 | ENLEALLVAL[K]K | 1xDSBSO_linker [K11] | P0A7L0 | 197 | A[K]PTQAK | 1xDSBSO_linker [K2] | P0A7L0 | 200 |
| y | n | 52.23 | Inter | 3 | GATGLGL[K]EAK | 1xDSBSO_linker [K8] | P0A7K2 | 82 | G[K]NGELTR | 1xDSBSO_linker [K2] | P0AG55 | 29 |
| y | n | 51 | Intra | 2 | YTAAITGAEG[K]IHR | 1xDSBSO_linker [K11] | P02358 | 35 | A[K]DER | 1xDSBSO_linker [K2] | P02358 | 106 |
| y | n | 51 | Intra | 1 | FPAIIYGG[K]EAPLAIELDHDK | 1xDSBSO_linker [K9] | P68919 | 34 | EI[K]VK | 1xDSBSO_linker [K3] | P68919 | 71 |
| y | n | 50.31 | Inter | 3 | SVAGF[K]IR | 1xDSBSO_linker [K6] | P62399 | 78 | [K]AGGSTR | 1xDSBSO_linker [K1] | P0A7L8 | 5 |
| y | n | 49.42 | Inter | 1 | A[K]LHDYYK | 1xDSBSO_linker [K2] | P62399 | 2 | VDRFN[K]R | 1xDSBSO_linker [K6] | P0A7M9 | 62 |
| y | n | 49.42 | Inter | 1 | AKDEADE[K]DAIATVNK | 1xDSBSO_linker [K8] | P0AG67 | 528 | [K]PELDAK | 1xDSBSO_linker [K1] | P0A7V3 | 108 |
| y | n | 49.37 | Intra | 1 | [K]GAIVTGK | 1xDSBSO_linker [K1] | P0AG67 | 450 | VEG[K]IK | 1xDSBSO_linker [K4] | P0AG67 | 370 |
| y | n | 49.21 | Inter | 2 | EFA[K]ANAK | 1xDSBSO_linker [K4] | P0A7J3 | 101 | VAVI[K]AVR | 1xDSBSO_linker [K5] | P0A7K2 | 71 |
| y | n | 47.63 | Inter | 2 | [K]SDQNVR | 1xDSBSO_linker [K1] | P0A7L0 | 54 | EA[K]IK | 1xDSBSO_linker [K3] | P0A7N9 | 53 |
| y | n | 47.4 | Inter | 1 | [K]AGFVTR | 1xDSBSO_linker [K1] | P0A7X3 | 100 | VL[K]FDR | 1xDSBSO_linker [K3] | P0AG67 | 247 |

| | | | | | | | | | | | | |
|---|---|---|---|---|---|---|---|---|---|---|---|---|
| y | n | 46.62 | Intra | 2 | [K]HNASR | 1xDSBSO_linker [K1] | P0A7U7 | 19 | TFI[K]K | 1xDSBSO_linker [K4] | P0A7U7 | 33 |
| y | n | 46.35 | Inter | 1 | EISMSI[K]R | 1xDSBSO_linker [K7] | P0A7S9 | 78 | GHAAD[K]K | 1xDSBSO_linker [K6] | P0A7U3 | 87 |
| y | n | 46.02 | Inter | 1 | [K]LQLVGVGYR | 1xDSBSO_linker [K1] | P0AG55 | 86 | AAGAN[K]VAVIK | 1xDSBSO_linker [K6] | P0A7K2 | 66 |
| y | n | 45.9 | Inter | 2 | VI[K]LDQK | 1xDSBSO_linker [K3] | P0AG67 | 158 | ASAV[K]R | 1xDSBSO_linker [K5] | P68679 | 54 |
| y | n | 44.12 | Inter | 1 | GHAAD[K]K | 1xDSBSO_linker [K6] | P0A7U3 | 87 | V[K]IGVAK | 1xDSBSO_linker [K2] | P0A832 | 126 |
| y | n | 43.63 | Intra | 1 | ELA[K]ASVSR | 1xDSBSO_linker [K4] | P0A7V3 | 49 | [K]GEDVEK | 1xDSBSO_linker [K1] | P0A7V3 | 80 |
| y | n | 42.39 | Inter | 1 | V[K]HPSEIVNVGDEITVK | 1xDSBSO_linker [K2] | P0AG67 | 229 | [K]AGFVTR | 1xDSBSO_linker [K1] | P0A7X3 | 100 |
| y | n | 41.94 | Intra | 2 | AAIEAAGG[K]IEE | 1xDSBSO_linker [K9] | P02413 | 141 | VT[K]GAR | 1xDSBSO_linker [K3] | P02413 | 129 |
| y | n | 41.82 | Intra | 2 | DHTLFA[K]ADGK | 1xDSBSO_linker [K7] | P0A7L8 | 62 | FEV[K]GPK | 1xDSBSO_linker [K4] | P0A7L8 | 72 |
| y | n | 40.95 | Inter | 1 | EA[K]DLVESAPAALK | 1xDSBSO_linker [K3] | P0A7K2 | 85 | G[K]NGELTR | 1xDSBSO_linker [K2] | P0AG55 | 29 |
